# Identification of a core transcriptional program driving the human renal mesenchymal-to-epithelial transition

**DOI:** 10.1101/2023.04.30.538857

**Authors:** John-Poul Ng-Blichfeldt, Benjamin J. Stewart, Menna R. Clatworthy, Julie M. Williams, Katja Röper

## Abstract

During kidney development, nephron epithelia arise *de novo* from fate-committed mesenchymal progenitors through a mesenchymal-to-epithelial transition (MET). Downstream of fate specification, transcriptional mechanisms that drive establishment of epithelial morphology through MET are poorly understood. We used human renal organoids derived from induced pluripotent stem cells, which recapitulate nephrogenesis, to investigate mechanisms controlling the renal MET programme. Multi-ome profiling of organoids revealed dynamic changes in gene expression and chromatin accessibility driven by transcriptional activators and repressors throughout renal MET. CRISPR-interference-based gene perturbation revealed that PAX8 is essential for initiation of MET in human renal organoids, contrary to mouse, by activating a cell adhesion programme. Furthermore, while Wnt/β-Catenin signalling specifies nephron fate, we find that it must be attenuated to allow HNF1B and TEAD transcription factors to drive completion of MET. These results reveal how the developing kidney balances fate-commitment and morphogenesis with implications for understanding both developmental kidney diseases and aberrant epithelial plasticity following adult renal tubular injury.

## Introduction

Emergence of complex tissues from simple tissue primordia begins with specification of tissue fate. Inductive signals, often produced by neighbouring tissues, activate gene regulatory networks within competent cells that establish identity and initiate cell differentiation and morphogenesis programmes. Ultimately, these lead to establishment of complex tissue shapes which enable specialised organ functions (Gilmour et al., 2017). The functional domains of most internal organs consist of epithelial sheets organised into tubular networks. Epithelial cells possess cell-scale morphological features that enable specialised tissue-scale functions, including apical-basal polarity, apical-lateral adherens junctions that provide adhesion and mechanical coupling to adjacent cells, and apical tight junctions that provide a membrane diffusion barrier and restrict paracellular flow of molecules (Rodriguez-Boulan & Macara, 2014). How tissue-specific fate programmes drive establishment of epithelial morphology is poorly understood.

The nephron, the functional unit of the mammalian kidney that filters the blood, is a convoluted tubular epithelial structure that arises through reciprocal inductive interactions between intermediate mesoderm-derived tissues during kidney development (Costantini & Kopan, 2010; McMahon, 2016). Nephrogenesis is initiated when the ureteric bud invades the metanephric mesenchyme. Inductive signals secreted by the metanephric mesenchyme cause branching morphogenesis of the ureteric bud epithelium, while in parallel, ureteric bud-secreted signals specify a nephron fate programme in neighbouring “cap” mesenchyme cells. This leads first to condensation of the cap mesenchyme into a pre-tubular aggregate, followed by a mesenchymal-to-epithelial transition (MET). The MET involves a stepwise assembly of intercellular junctions and *de novo* establishment of apical-basal polarity to give rise to the renal vesicle, the first polarised epithelial precursor of the nephron. This renal vesicle subsequently elongates into the comma-and then S-shaped body and ultimately fuses with the ureteric bud tip to form the nascent nephron (Costantini & Kopan, 2010; McMahon, 2016). Nephrogenesis occurs entirely prenatally and nephron endowment at birth is an important determinant of adult kidney health, with low nephron number predisposing to a range of adult kidney diseases (Charlton et al., 2021). Therefore, a complete knowledge of mechanisms controlling nephron development is critical to better understand and prevent diseases of renal insufficiency.

The primary nephrogenic inductive cue secreted by the ureteric bud is Wnt9b (Carroll et al., 2005), and activation of Wnt/β-Catenin signalling downstream of Wnt9b is necessary and sufficient to specify the nephron fate programme required for renal MET (Park et al., 2007). However, sustained Wnt/β-Catenin signalling appears to inhibit completion of renal MET (Park et al., 2012; Park et al., 2007), and Wnt/β-Catenin signalling has been reported to be able to drive the opposite process, an epithelial-to-mesenchymal transition (EMT), in various instances including gastrulation and in cancer [e.g. (Kemler et al., 2004; Yook et al., 2005; Yook et al., 2006) or reviews (Gonzalez & Medici, 2014; Ng-Blichfeldt & Röper, 2021; Pei et al., 2019)]. Therefore, how activation of the nephron fate programme by Wnt/β-Catenin signalling leads to establishment of epithelial morphology during nephrogenesis is unclear.

Most insights into mechanisms of nephrogenesis derive from mouse models, however, recent studies highlight divergence between developing mouse and human kidneys, particularly in patterns of transcription factor expression during nephrogenesis (Lindstrom, Guo, et al., 2018; Lindstrom, McMahon, et al., 2018; Lindstrom, Tran, et al., 2018). Transcriptome profiling at single cell resolution is emerging as a powerful tool for elucidating gene expression programmes of human kidney development where functional studies are not possible (Hochane et al., 2019; Lindstrom, Guo, et al., 2018; Schreibing & Kramann, 2022; Stewart et al., 2019), however, how these programmes are controlled by transcription factors remains largely unexplored. The recent development of a “multi-omic” approach in which transcriptome profiling is combined with chromatin accessibility profiling has enabled insights into how combinations of transcription factors drive gene expression programmes (Granja et al., 2021). Moreover, the advent of human induced pluripotent stem cell (iPSC)-derived renal organoids which recapitulate nephrogenesis, now allows for the functional investigation of human-specific mechanisms of kidney development (Kumar et al., 2019; Nishinakamura, 2019; Takasato et al., 2015).

To investigate transcription factor regulation of the earliest morphogenetic changes in the developing human kidney, we applied paired transcriptome and chromatin accessibility profiling to human iPSC-derived renal organoids across the time period of renal MET combined with *in vivo/organoid* analyses and functional studies. We identify dynamic gene expression and chromatin accessibility signatures driven by both transcriptional activators and repressors during human renal MET, and identify PAX8 as a critical upstream regulator of human renal MET. Whereas it was previously shown that mice lacking PAX8 have no kidney defects (Bouchard et al., 2002), we find using CRISPR-interference based gene perturbation that PAX8 is essential for initiation of MET in human renal organoids. We further find that attenuation of Wnt/β-Catenin signalling is required for HNF1B and TEAD family transcription factor activity to drive completion of MET. Our study reveals how the developing kidney balances fate-commitment and morphogenesis, whilst providing a rich resource for the community to now explore early human kidney morphogenesis and development in greater depth.

## Results

### Human iPSC-derived kidney organoids recapitulate renal mesenchymal-to-epithelial transition

We set out to investigate how a key step in nephron development, the establishment of epithelial polarity preceding the formation of the renal vesicle, is induced and controlled during human nephrogenesis. We adapted a published protocol to derive renal organoids from human iPSCs (Kumar et al., 2019; Takasato et al., 2015) (Fig. 1A), whereby nephron fate is induced through Wnt/β-Catenin pathway activation with the GSK3 inhibitor CHIR99021 (CHIR; (Kumar et al., 2019; Takasato et al., 2015)) combined with suspension culture of spherical organoids. Analysis of epithelial polarity markers atypical protein kinase C (aPKC; apical) and integrin β1 (ITGB1; basal), and epithelial junctional markers zonula occludens-1 (ZO1; tight junctions) and E-Cadherin (CDH1; adherens junctions), showed that the MET takes place between day 10 and 14 of the protocol (Fig. 1A-C). All markers became progressively more localised along the apical-basal axis, resembling what has been described in the developing kidney *in vivo* (Horster et al., 1999). Organoids grown to day 24 of the protocol contained recognisable renal structures that showed differentiation into podocytes (WT1^+^), proximal tube cells (LTL^+^, ECAD^+^) and distal tube cells (LTL^-^, ECAD^+^) (Supplemental Fig. S1B). Thus, human renal organoids generated with our protocol recapitulate nephrogenesis and contain expected nephron cell types.

**Figure 1.**
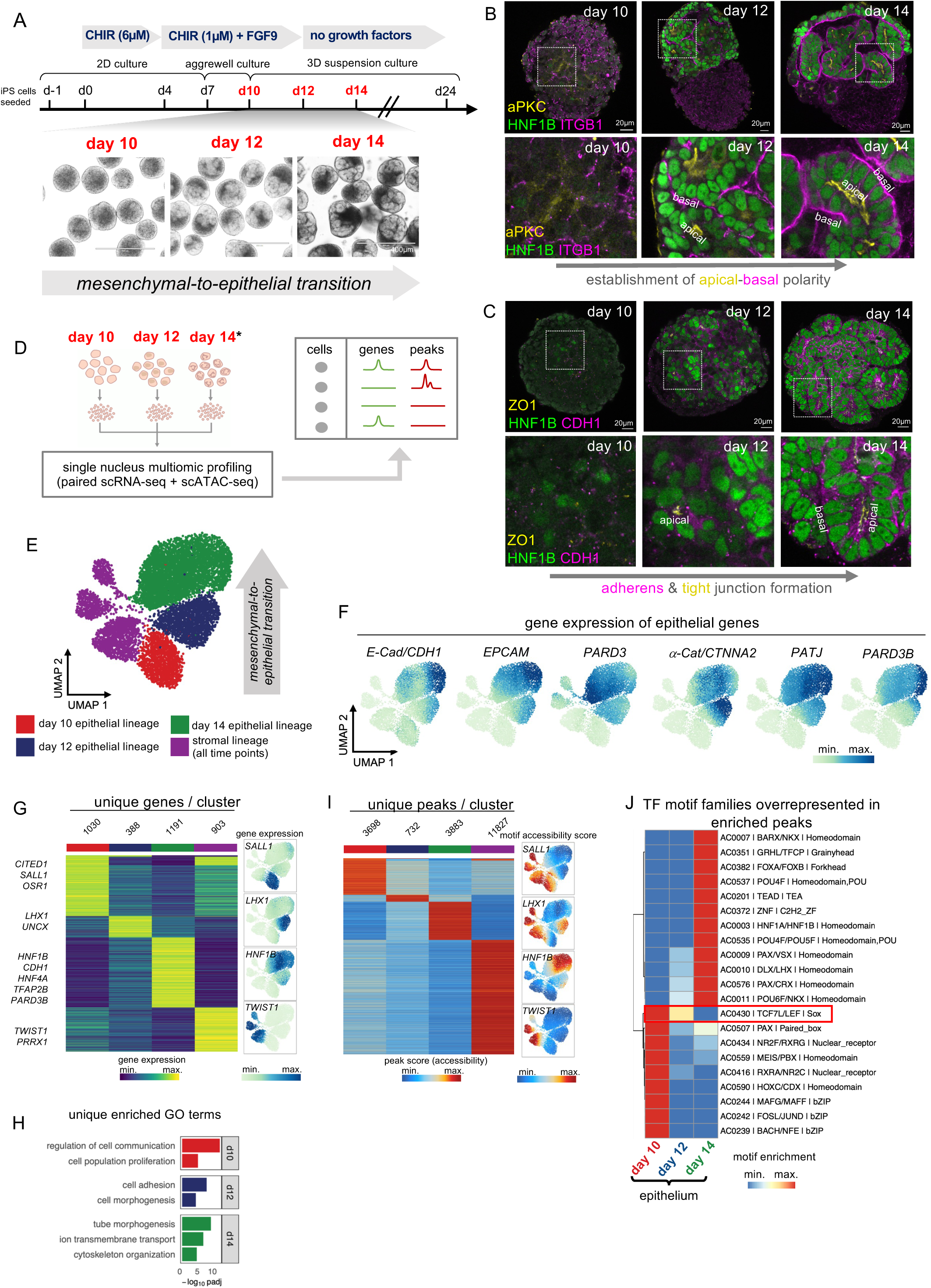
RNA-and ATAC-seq profiling of human renal organoids reveals dynamic gene expression and chromatin accessibility signatures during renal MET. (A) Schematic of protocol used to generate human renal organoids from iPSCs, with corresponding light microscopy images of organoids during MET (day 10-14) (adapted from (Kumar et al., 2019; Takasato et al., 2015)). Scale bars 400µm. (B, C) Immunofluorescence showing expression of (B) aPKC (yellow), integrin β1/ITGB1 (magenta), and HNF1B (green), or (C) ZO-1 (yellow), E-Cadherin/CDH1 (magenta), and HNF1B (green), in renal organoids over the time course of MET. Dotted white boxes indicate positions of magnification panels below, scale bars are 20µm. (D) Schematic of renal organoid multi-ome profiling strategy for batch 2 and batch 3 individually. *For batch 3, two day 14 samples were sequenced in parallel: one treated with DMSO from day 12 to 14, and one treated with 3µM CHIR from day 12 to 14, relevant to Fig. 6 (see also Supplemental Fig. 11E). (E) UMAP representation of 9,147 cells from multi-ome batch 3 projected according to ATAC data, clustered by epithelial lineage time point, and stromal lineage (all time points combined). (F) Gene expression levels of *CDH1, EPCAM, PARD3, CTNNA2, PATJ,* and *PARD3B* overlaid on UMAP plots. (G) Heat map of expression of marker genes for clusters in (E) determined by snRNA-seq with representative markers annotated (left, log2FC > 1, FDR < 0.01, two-sided Wilcoxon rank-sum test; log2FC: log2 fold change; FDR: false discovery rate). Gene expression UMAP plots of cluster-specific transcription factors (right). (H) Unique GO terms represented in enriched genes from (G), full list in Supplemental Table 3. (I) Heat map of marker peaks of accessible chromatin for clusters in (E) determined by snATAC-seq (left, log2FC > 1, FDR < 0.01, two-sided Wilcoxon rank-sum test). Motif accessibility UMAP plots of cluster-specific transcription factors from (G) (right). (J) Heat map of transcription factor motif archetypes (Vierstra et al., 2020) enriched in epithelial-specific peaks from (I). Annotated are archetype codes, archetypal transcription factors and DNA-binding class. Wnt/β-Catenin transcriptional effectors TCF7/LEF are highlighted (red). See also Supplemental Figs S1 and S2.

To characterise the renal organoids around the time frame of epithelial polarisation in depth, we performed single cell RNA-sequencing (scRNA-seq) and single nucleus multi-ome sequencing (paired snRNA-seq and snATAC-seq) on organoids from four time points (day 10, day 12, day 14 and day 24) across three batches (for details see STAR METHODS). First, scRNA-seq data and snRNA-seq data (without corresponding snATAC-data) were pooled, which after filtering resulted in 23,856 cells that separated into 2 clusters when visualised as a uniform manifold approximation projection (UMAP) with coarse clustering (Supplemental Fig. S1F-H, see STAR METHODS). One cluster was enriched for *PAX8, PAX2* and epithelial cell adhesion molecule (*EPCAM*) gene expression, and so was annotated as the epithelial lineage, whereas the other cluster was enriched for *PRRX1, TWIST1* and *COL1A2* gene expression, and so was annotated as the stromal lineage (Supplemental Fig. S1F-H). Isolation and re-clustering of the epithelial lineage resulted in 15,917 cells which were annotated according to time point, cell cycle phase, and expression of known nephron markers (Supplemental Fig. S1I-L) (Lindstrom et al., 2021). This revealed the presence of nephron progenitors and differentiating tube epithelial cells, proliferating cells, as well as differentiated podocytes (*MAFB*^+^), proximal tube (*HNF4A*^+^), distal tube (*TFAP2B*^+^) and connecting tube (*GATA3*^+^) cells (Supplemental Fig. S1J), consistent with human renal organoids recapitulating human nephrogenesis.

At day 10, most genes characteristic of the epithelial state were lowly expressed, consistent with cells still being mainly mesenchymal, whereas at day 14, *EPCAM*, E-Cadherin/*CDH1* and the polarity component partitioning defective 3B (*PARD3B*) were upregulated and showed maximal expression (Supplemental Fig. S1M), consistent with completion of MET by day 14. Tight junction components Claudin 7 (*CLDN7*), Claudin 3 (*CLDN3*) and the apical polarity determinant Crumbs 3 (*CRB3*) were upregulated at day 14 and further upregulated at day 24, suggesting progressive maturation of epithelial cells (Supplemental Fig. S1M). Expression of markers of differentiated nephron cell types, such as *MAFB*, *HNF4A* and *TFAP2B*, was highest at day 24, consistent with this time point representing a fully differentiated nephron (Supplemental Fig. S1N).

To investigate the relevance of renal organoid morphogenesis to human fetal kidney development *in vivo*, we analysed our dataset alongside a previously described scRNA-sequencing dataset of 6 human fetal kidney samples collected between post conception weeks 7 and 16 (Stewart et al., 2019). This dataset consisted of 17,994 cells, and segregated into epithelial (*PAX8*^+^) and stromal (*TWIST1*^+^) lineages (Supplemental Fig. S2A and B). Projection of organoid scRNA-seq data onto the human fetal dataset by time point revealed congruence between iPSC-derived renal organoids and fetal kidney, with acquisition of differentiated cell types (podocytes, proximal tube and distal tube epithelial cells) at day 24 (Supplemental Fig. S2A-C), indicating that renal organoids represent an experimentally tractable model system that faithfully recapitulates human renal MET *in vivo*.

### Multi-ome profiling of human renal organoids reveals dynamic gene expression and chromatin accessibility signatures during renal MET

To gain insights into transcriptional control of renal MET and early nephron morphogenesis, we next interrogated the multi-ome sequencing dataset of human renal organoids to determine gene expression profiles and genome wide chromatin accessibility at single cell resolution. We focused on early time points with higher temporal resolution, namely, days 10, 12 and 14 from two batches (batches 2 and 3), to capture the time period of renal MET; both batches revealed similar results, and data from batch 3 are shown unless otherwise indicated (for details see STAR METHODS). Filtering of cells from batch 3 resulted in 9,147 cells that separated into epithelial (*PAX8*^+^) and stromal (*TWIST1*^+^) lineages upon coarse clustering (Supplemental Figure S3A and B, Fig. 1E). Gene expression of *EPCAM*, E-Cadherin/*CDH1*, and α-Catenin (*CTNNA2*), and the polarity components *PATJ*, *PARD3* and *PARD3B* increased within the epithelial lineage from day 10 to day 14, consistent with the progression of epithelial polarisation and MET (Fig. 1F). Higher resolution clustering revealed that the epithelial lineage at day 14 contained discrete populations of cells expressing *WT1*, *HNF4A* or *TFAP2B,* corresponding to podocytes and proximal and distal nephron, respectively, indicating that nephron segment identity is specified within renal organoids by day 14 concomitant with MET (Supplemental Fig. S3C-G; Supplemental Table 1).

We observed dynamic gene expression of established nephron markers within the epithelial lineage throughout MET. *CITED1, SALL1,* and *OSR1* were enriched at day 10, consistent with a mesenchymal nephron progenitor state, whereas the renal vesicle marker *LHX1* was enriched at day 12. The epithelial markers *HNF1B, CDH1, PARD3B,* together with *HNF4A* and *TFAP2B* which control proximal tube and distal tube differentiation, respectively (Marable et al., 2020; Marneros, 2020), were enriched at day 14, indicating that renal organoids follow the transcriptional trajectory of human nephrogenesis (Lindstrom et al., 2021) (Fig. 1K; Supplemental Table 2).

Gene ontology analysis (Fig. 1H; Supplemental Table 3) revealed enrichment at day 10 for cell-cell communication and proliferation, consistent with the rapid increase in size of organoids around this time point, whereas day 12 showed enrichment of cell adhesion and cell morphogenesis genes, consistent with expression of adherens and tight junction components (Fig. 1C and F). Day 14 showed enrichment of genes involved in cytoskeleton organisation and tube morphogenesis, consistent with the proper organisation of adherens and tight junction complexes and establishment of tubular morphology (Fig. 1C). Ion transport components were also overrepresented at day 14, suggesting initial stages of nephron segmentation. Thus, gene expression dynamics in human renal organoids between days 10 and 14 reveal dynamic cell-and tissue-level morphogenetic processes driving the renal MET, and suggest that expression of adherens and tight junction components are one of the earliest events in MET, preceding both junctional organisation and nephron segment differentiation.

The analysis of ATAC data revealed dynamic chromatin accessibility profiles throughout renal MET (Fig. 1I). These data were analysed for transcription factor binding motif archetypes to identify candidate transcription factors driving these changes (Vierstra et al., 2020), which revealed two classes of motif archetypes (Fig. 1J). The first class was enriched at day 10 and decreased by day 14, which included motifs for Wnt/β-Catenin transcriptional effectors TCF7/LEF, indicating initial high Wnt/β-Catenin signalling followed by later attenuation (Fig. 1J). The second class of motif archetypes was absent at day 10 and increased by day 14, which included motifs for Grainyhead-like (GRHL) factors which have been suggested to drive MET in other systems (Ng-Blichfeldt & Röper, 2021), tea-domain containing (TEAD) factors which are downstream effectors of the Hippo/Yes-associated protein (YAP) signalling pathway (Zhao et al., 2008), and the renal vesicle stage markers HNF1A/B (Lindstrom, Tran, et al., 2018). From these data we propose a model whereby renal MET is biphasic, with Wnt/μ-Catenin signalling being initially high and becoming attenuated as MET progresses, which in turn permits multiple transcription factors to drive completion of renal MET.

### Comparison of epithelium and stroma reveals early drivers of renal MET

We next asked which transcription factors were central to initiating MET by investigating transcription factors that best defined the epithelial lineage compared to the stromal lineage. Coarse clustering of multi-ome cells resulted in 2 clusters corresponding to the epithelial (*PAX8*^+^) and stromal (*TWIST1*^+^) lineages (Fig. 2A-C; Supplemental Table 4).

**Figure 2.**
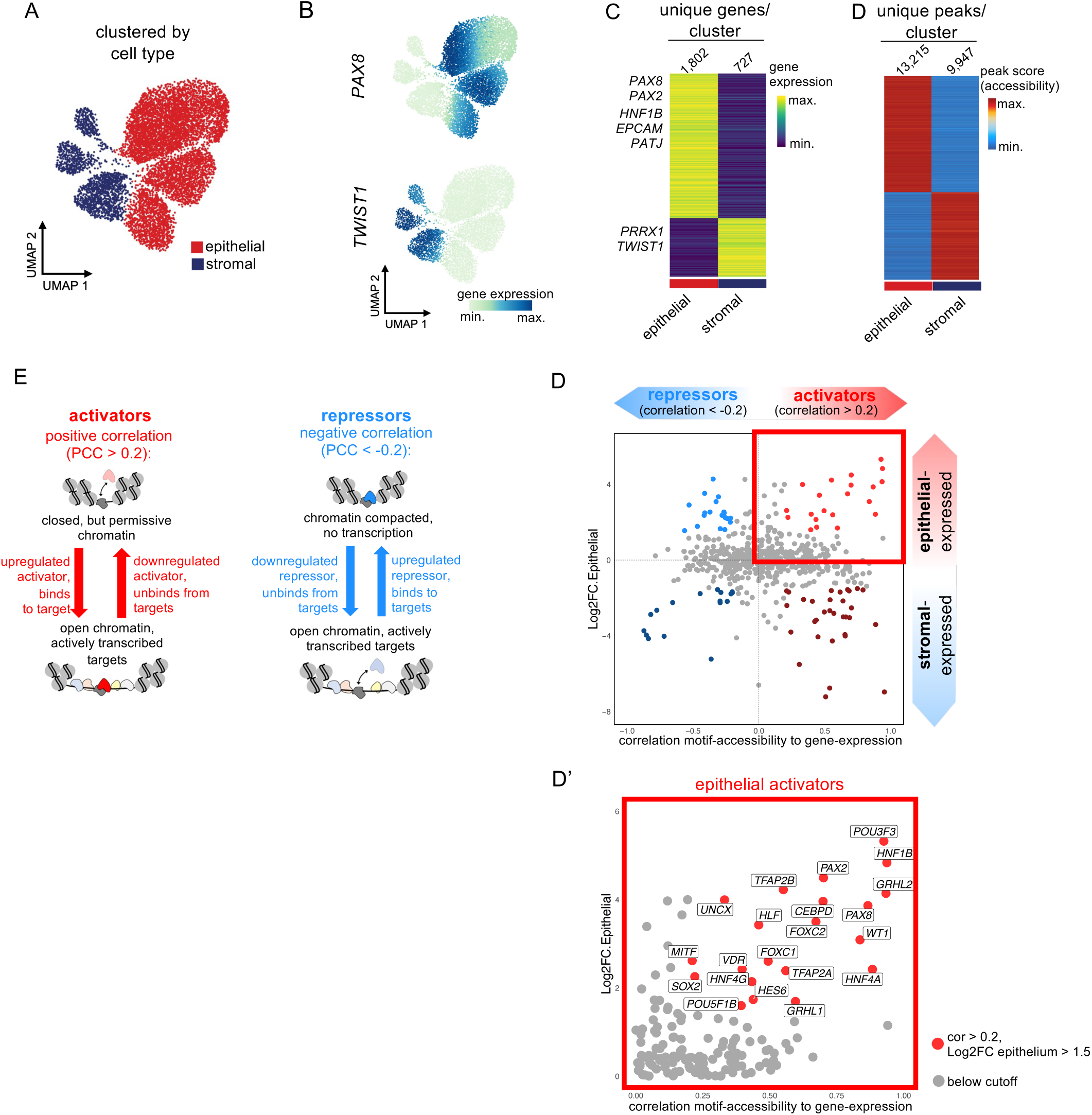
Early transcriptional activators driving renal mesenchymal-to-epithelial transition. (A) UMAP of 9,147 cells from multi-ome batch 3 projected according to ATAC data, with clusters annotated according to epithelial or stromal lineage. (B) UMAP plots of multi-ome cells from batch 3, coloured by *PAX8* gene expression (top) and *TWIST1* gene expression (bottom). (C) Heat map of expression of marker genes for clusters in (A) determined by snRNA-seq with representative marker genes annotated (log2 FC > 1, FDR < 0.01, two-sided Wilcoxon rank-sum test). (D) Heat map of marker peaks of accessible chromatin for clusters in (A) determined by snATAC-seq (left, log2 FC > 1, FDR < 0.01, two-sided Wilcoxon rank-sum test). (E) Schematic of the principle by which correlation values, between transcription factor gene expression and corresponding motif accessibility, were used to classify transcription factors as either transcriptional repressors or activators. PCC, Pearson correlation coefficient. (F) Transcription factors plotted according to PCC of gene expression versus corresponding motif accessibility, and log2FC gene expression in the epithelial compared to the stromal lineage as determined by snRNA-seq. Thresholds used to colour points according to principle outlined in (D): PCC > 0.2, Log2FC epithelial lineage > 1.5 (red = activators), PCC < -0.2, Log2FC epithelial lineage > 1.5 (blue = repressors). See also Supplemental Figs S3 and S4.

Analysis of ATAC data revealed differentially accessible chromatin regions between clusters (Fig. 2D). To investigate transcription factors regulating the epithelial state, correlations were derived between transcription factor gene expression and accessibility of corresponding motifs across all cells (Fig. 2E, see STAR METHODS). A positive correlation implies transcription factor binding at target loci and recruitment of epigenetic modifiers that drive chromatin opening and increased motif accessibility, a characteristic of transcriptional activators. A negative correlation implies recruitment of chromatin compactors that cause a reduction in motif accessibility, a characteristic of transcriptional repressors. This analysis revealed transcriptional activators and repressors selectively expressed in either epithelial or stromal lineages (Fig. 2F). Corresponding analysis of batch 2 revealed similar results (Supplemental Fig. S4A-E). Among top epithelial activators were transcription factors that show segment-restricted expression within the mature nephron (*TFAP2B*, *HNF4A*), and *PAX2* and *PAX8*, which are expressed prior to acquisition of segment identity, and thus are candidate novel upstream regulators of MET.

### PAX8 expression precedes induction of MET in renal organoids and in human fetal kidney *in vivo*

PAX2 and PAX8 are closely related paired box transcription factors that are both expressed in developing human and mouse kidneys (Lindstrom, Tran, et al., 2018). We first analysed a published scRNA-seq dataset of human fetal kidney (Stewart et al., 2019), specifically focussing on 8,862 cells of the fetal nephron lineage, to determine *PAX2* and *PAX8* gene expression in the developing human nephron *in vivo*. *PAX2* was expressed in uncommitted (*SIX1^+^/SIX2^+^/EYA1^+^*) cap mesenchyme, and also in committed cap mesenchyme and subsequent committed epithelium (corresponding to renal vesicle and comma-/S-shaped body stage cells; Fig. 3A-C, Supplemental Fig. S2D), consistent with published immunofluorescence analysis of PAX2 in human fetal kidney sections (Lindstrom, Tran, et al., 2018). Notably, expression of *PAX2* already in uncommitted cap mesenchyme indicates that PAX2 alone is insufficient to initiate renal MET. By contrast, in the scRNA-seq dataset of *in vivo* human fetal kidney, PAX8 was absent from uncommitted cap mesenchyme, and instead was restricted to committed cap mesenchyme and subsequent committed epithelium (Fig. 3A-C, Supplemental Fig. S2D), an expression pattern seen in published immunofluorescence analysis of PAX8 in human fetal kidney sections (Lindstrom, Tran, et al., 2018), and also seen in mice (Carroll et al., 2005; Plachov et al., 1990); this pattern is highly suggestive of PAX8 having a central role in establishing the renal MET programme.

**Figure 3.**
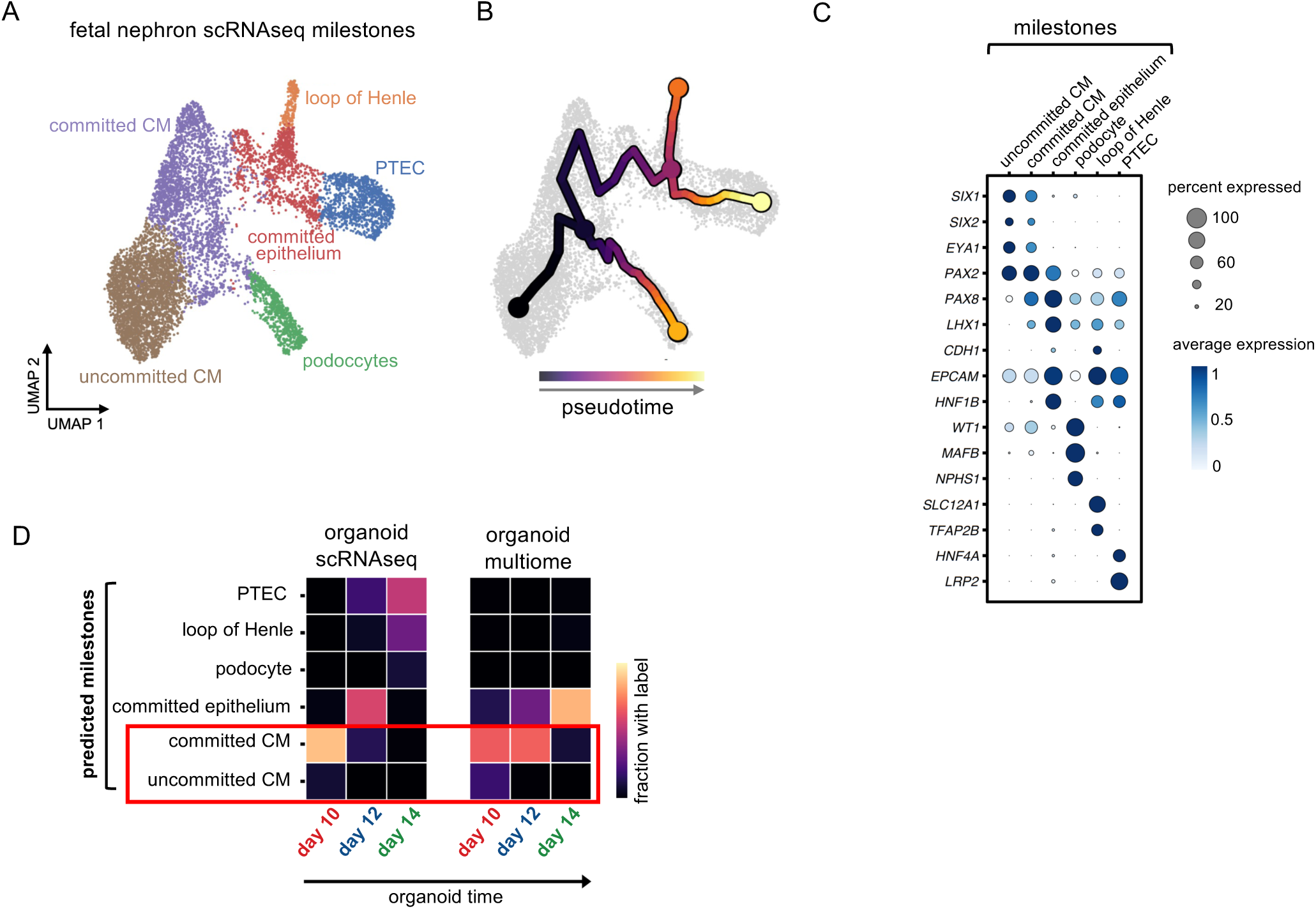
Human iPSC-derived kidney organoids show congruence with committed cap mesenchyme and subsequent epithelial derivatives in human fetal kidney *in vivo*. (A) UMAP plot of 8,862 sc-RNA-seq cells from 6 human fetal kidney samples harvested between post conception week (pcw) 7 and 16, showing nephron epithelial lineage only (Stewart et al., 2019), with developmental milestones annotated. (B) UMAP plot of 8,862 sc-RNA-seq cells as in (A) with nephron pseudotime trajectory overlaid. (C) Expression of marker genes of nephron developmental stages within fetal sc-RNA-seq data. (D) Prediction of milestone labels within organoid scRNA-seq and snRNA-seq data. See also Supplemental Fig. S2

Comparing the transcriptomes of renal organoids and fetal kidney revealed that organoids at day 10 showed closest alignment with committed cap mesenchyme *in vivo* (Fig. 3D), consistent with organoids specifically and faithfully recapitulating induction of cap mesenchyme and subsequent MET. Interrogating the multi-ome data, we found that *PAX8* showed high gene expression and high motif accessibility at day 10, with gene expression slightly reduced at day 14 (Supplemental Fig. S5A), mainly retained in podocytes (Supplemental Fig. S3C-G). *PAX2* showed highest gene expression at day 12, but high accessibility of the corresponding motif already at day 10 (Supplemental Fig. S5A); PAX2 and PAX8 share a similar DNA recognition motif (Phelps & Dressler, 1996), and therefore accessibility of the PAX2 motif at day 10 was likely due to high *PAX8* expression at this time point. The earlier expression of PAX8 compared to PAX2 in organoids was confirmed by immunofluorescence (Supplemental Fig. S5B). The renal organoid model therefore enables the opportunity to disentangle the precise role of PAX8 from PAX2 in committed nephrogenic mesenchyme, which is usually masked *in vivo* by persistence of PAX2 from uncommitted cap mesenchyme and potential partial redundancy between PAX8 and PAX2 (Bouchard et al., 2002; Narlis et al., 2007).

### PAX8 is required for the initiation of MET in human renal organoids

The above data support the hypothesis that PAX8 is a critical upstream regulator of human renal MET. To test this, dead-Cas9-KRAB-based CRISPR interference (Tang et al., 2022) was used to generate iPSCs that allow suppression of PAX8 gene expression under the control of doxycycline (Dox), with cytoplasmic GFP as a readout for the response to Dox (this cell line is hereafter referred to as PAX8-dCas9-KRAB; Fig. 4A; see STAR METHODS). As the *PAX8* gene was expressed from day 7 in the organoid protocol (Supplemental Fig. S5C), PAX8-dCas9-KRAB iPSCs differentiated into organoids were treated with 1µM Dox from day 4, which led to ∼80% knockdown of *PAX8* gene expression compared to untreated control organoids at day 14 (*** p < 0.001; Fig. 4B). *PAX8* knockdown led to a significant corresponding reduction in *CDH1* gene expression at day 14 (∼80%, * p < 0.05; Figure 4C). Control organoids at day 14 did not show GFP by immunofluorescence, indicating lack of knockdown cassette activity (Fig. 4D), and contained cells that exhibited PAX8 staining that also expressed E-Cadherin as we observed previously (Fig. 4D). By contrast, Dox-treated organoids at day 14 showed heterogeneous GFP expression, and strongly reduced PAX8 staining. Cells that retained PAX8 also lacked GFP, and thus were not responding to Dox (Fig. 4D, E). In these organoids, E-Cadherin expression was mostly depleted, and where present, was confined to the PAX8^+^ GFP-lacking cells (Fig. 4D). Quantification revealed ∼80% reduction in E-Cadherin signal volume and hence in the degree of epithelialisation after PAX8-knockdown compared to control organoids (*** p< 0.0001; Fig. 4E).

**Figure 4.**
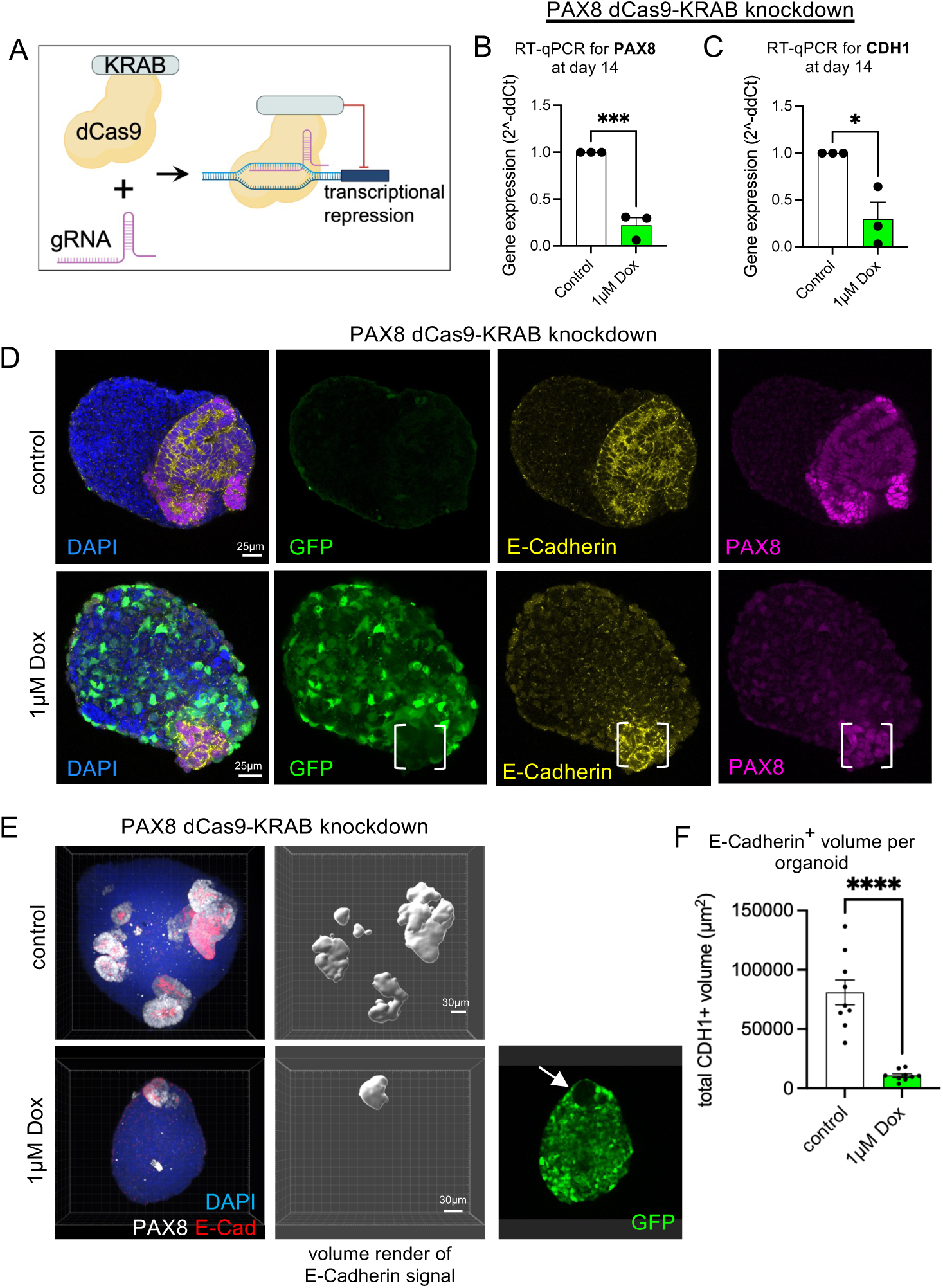
PAX8 is a critical upstream regulator of MET in human renal organoids. (A) Schematic of the dCas9-KRAB CRISPR interference gene perturbation system (Tang et al., 2022). (B, C) RT-qPCR for (B) *PAX8* and (C) *E-Cadherin/CDH1* in organoids generated from PAX8-dCas9-KRAB iPSCs and harvested at day 14, either with no treatment (control, white bars) or following Dox treatment from day 4 (1μM Dox, green bars). N = 3 independent organoid batches. Dots represent data from each batch normalised to corresponding control, with bars representing mean ± SEM. Unpaired t-test, * (p < 0.05), *** (p < 0.001), compared to corresponding control. (D) Immunofluorescence of organoids generated from PAX8-dCas9-KRAB iPSCs and harvested at day 14, either with no treatment (control, top) or following Dox treatment from day 4 (1μM Dox, bottom), showing GFP (green), PAX8 (magenta), E-Cadherin/CDH1 (yellow) with DAPI as counterstain (blue, nuclei). White brackets indicate PAX8^+^CDH1^+^ cells confined to a GFP^-^ region. Scale bars 25µm. (E) Immunofluorescence images showing projected z-stacks of whole organoids of PAX8-dCas9-KRAB organoids at day 14, either with no treatment (control, top) or following Dox treatment from day 4 (1 μM Dox, bottom), showing PAX8 (white), E-Cadherin (red), and DAPI (blue, nuclei) alongside volume render of corresponding E-Cadherin/CDH1 signal (middle), and GFP expression after Dox treatment (green, right bottom). Arrow points to GFP^-^ region that contains PAX8^+^CDH1^+^ cells. Scale bars 30µm. (F) Quantification of volume-rendered E-Cadherin/CDH1 signal in PAX8-dCas9-KRAB organoids at day 14 either with no treatment (Control, white bar) or following Dox treatment from day 4 (1μM Dox, green bar). n = 9 organoids across N = 2 independent batches. Dots represent data from individual organoids, with bars representing mean ± SEM. Unpaired t-test, **** (p < 0.0001). See also Supplemental Fig. S5.

We next assessed the role of PAX2 in human renal MET by generating an iPSC line that allowed Dox-dependent suppression of PAX2 expression (referred to as PAX2-dCas9-KRAB, see STAR METHODS). When differentiated to organoids, treatment with 1µM Dox from day 4 resulted in ∼80% knockdown of *PAX2* gene expression compared to untreated control organoids (*** p < 0.001, unpaired t-test; Supplemental Fig. S5D). At day 14, control organoids lacked GFP and contained tubular structures positive for PAX2 and E-Cadherin (Supplemental Fig. S5E). Dox-treated organoids showed heterogeneous GFP expression but contained tubular structures positive for PAX2 and GFP in a mutually exclusive pattern, indicating that GFP^+^ cells successfully knocked down PAX2 expression (Supplemental Fig. S5E). GFP^+^ PAX2^-^ cells expressed E-Cadherin and contributed to tubular structures, indicating that PAX2 is dispensable for generation of nephron epithelium from human iPSCs, in accordance with a previous report (Kaku et al., 2017).

Our analyses in human renal organoids thus reveal a hitherto unappreciated central role for PAX8 in initiating MET during human kidney development that, in contrast to mouse kidney development *in vivo* (Bouchard et al., 2002; Narlis et al., 2007), cannot be compensated for by PAX2.

### PAX8 activates a cell adhesion gene programme to initiate MET

We next investigated the genetic programme activated downstream of PAX8 that drives MET. In day 10 organoids, PAX8 localised to cells that coalesced in the centre of the organoids, analogous to the pre-tubular aggregate; moreover, PAX8^+^ cells selectively expressed HNF1B and E-Cadherin at day 12 (Fig. 5A). Footprint analysis of ATAC data revealed occupancy at both PAX8 and PAX2 motifs in the epithelial lineage at all time points, but not in the stromal lineage (Fig. 5B), suggesting that PAX8 drives expression of target genes throughout MET. Next, the PAX8 regulon was determined by analysis of peak to gene links (Supplemental Fig. 5F, STAR METHODS). Gene ontology analysis of the PAX8 regulon revealed enrichment for cell adhesion and cadherin binding (Fig. 5C, Supplemental Table 5). Interestingly, the PAX8 regulon contained Cadherin-6 (*CDH6*), which is one of the earliest cadherins expressed in the mouse condensed mesenchyme *in vivo* (Mah et al., 2000) and was expressed in renal organoids at day 10 prior to other cadherins (Supplemental Fig. S5G). Examination of the *CDH6* genomic locus revealed 4 accessible peaks significantly linked to *CDH6* (PCC < 0.4, FDR <0.0001; Fig. 5D). One peak 86 kb upstream of the *CDH6* TSS which was accessible in the epithelial lineage throughout MET, but not in the stroma, showed enrichment for PAX8 motifs, consistent with PAX8 driving expression of *CDH6* by directly binding to a regulatory element within this genomic region (Fig. 5D). In support of this, PAX8^+^ cells also expressed CDH6 in day 10 organoids (Supplemental Fig. S5H).

**Figure 5.**
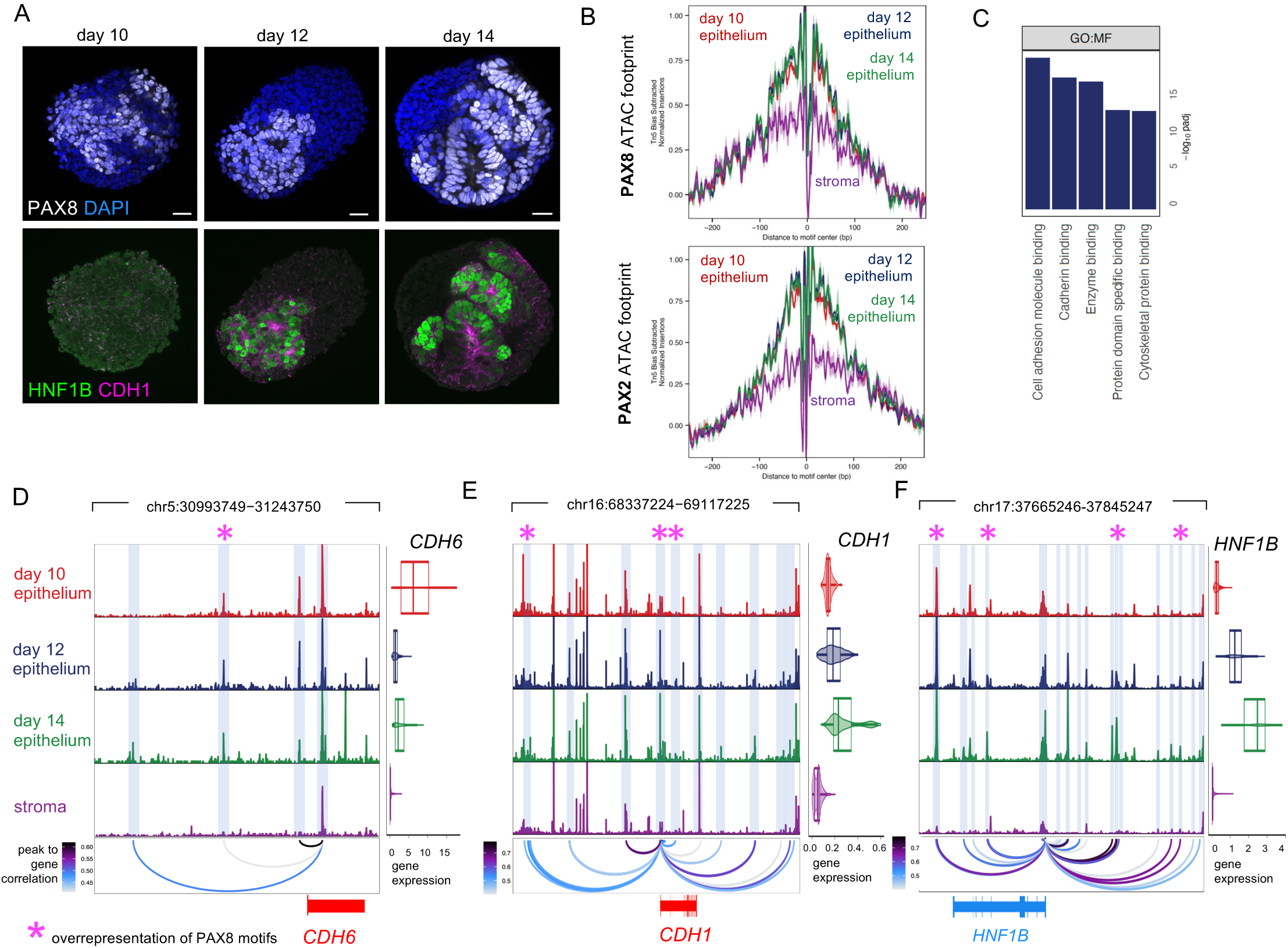
PAX8 is activates a cell adhesion gene expression programme to initiate MET in human renal organoids. (A) Immunofluorescence analysis of organoids showing PAX8 (white, top) with DAPI as counterstain (blue, nuclei), with corresponding expression of HNF1B and E-Cadherin/CDH1 (green and magenta, respectively, bottom) at day 10, 12 and 14. Scale bars 25µm. (B) Footprint analysis of PAX8 and PAX2 motifs using cluster annotations from (Fig. 1E). (C) GO terms overrepresented in the PAX8 regulon (full list in Supplemental Table 5). (D-F) Chromatin accessibility tracks for (D) *CDH6*, (E) *E-Cadherin/CDH1* and (F) *HNF1B* loci, split according to clusters annotated in (Fig. 1E), epithelium day 10, 12, 14 and stroma (all time points combined), with peak to gene links highlighted. Violin plots of gene expression determined by snRNA-seq (box center line, median; limits, upper and lower quartiles; whiskers, 1.5x interquartile range). Purple asterisks indicate overrepresentation of PAX8 motifs as determined by analysis with HOMER (see STAR METHODS). See also Supplemental Fig. S5.

*CDH1* and *HNF1B* were also contained within the PAX8 regulon and therefore potentially regulated by PAX8, consistent with the presence of E-Cadherin and HNF1B protein in PAX8^+^ cells in day 12 organoids (Fig. 5A). Analysis of the *CDH1* genomic locus revealed 25 peaks significantly linked to *CDH1* (PCC < 0.4, FDR <0.0001; Fig. 5E), including two peaks corresponding to enhancers within the second intron of *CDH1* (Alotaibi et al., 2015). PAX8 motifs were overrepresented in three *CDH1*-linked peaks. Analysis of the *HNF1B* locus revealed 24 peaks significantly linked to *HNF1B* (PCC < 0.4, FDR <0.0001; Fig. 5F); PAX8 motifs were overrepresented in 3 of these peaks.

We therefore propose that PAX8 is a critical component of the Wnt-dependent nephron fate programme, and that PAX8 drives human renal MET by directly activating expression of genes that establish epithelial polarity and intercellular junctions. Altogether, these data help to explain how establishment of the nephron fate programme drives the emergence of complex tissue morphology during kidney development.

### A combination of transcriptional activators and repressors drives human renal MET

The above analysis places PAX8 as a critical upstream regulator that initiates MET in human renal development, likely directly downstream of Wnt/β-Catenin signalling (Schmidt-Ott et al., 2007). We next interrogated the multi-ome dataset to identify transcription factors that regulate the completion of polarisation and epithelialisation during MET. Using pseudotime analysis, a trajectory representing the emergence of renal tube epithelial cells via MET in kidney organoids was inferred (Fig. 6A). Gene expression of *EPCAM, E-Cadherin/CDH1* and *PARD3B* increased along organoid tube pseudotime consistent with progression of MET (Fig. 6B). Comparison with the fetal nephron pseudotime trajectory revealed that the organoid tube pseudotime trajectory aligned with the transition from committed cap mesenchyme to committed epithelium, and thus corresponded to the period of MET *in vivo* (Fig. 3D and Fig. 6C). By correlating transcription factor gene expression and motif accessibility along pseudotime, 52 activators and 41 repressors were identified that showed dynamic changes along MET (FDR < 0.001; Fig. 6D and E); these were ordered by gene expression along pseudotime to reveal a temporal sequence of activator and repressor function during renal MET (Fig. 6D and E).

**Figure 6.**
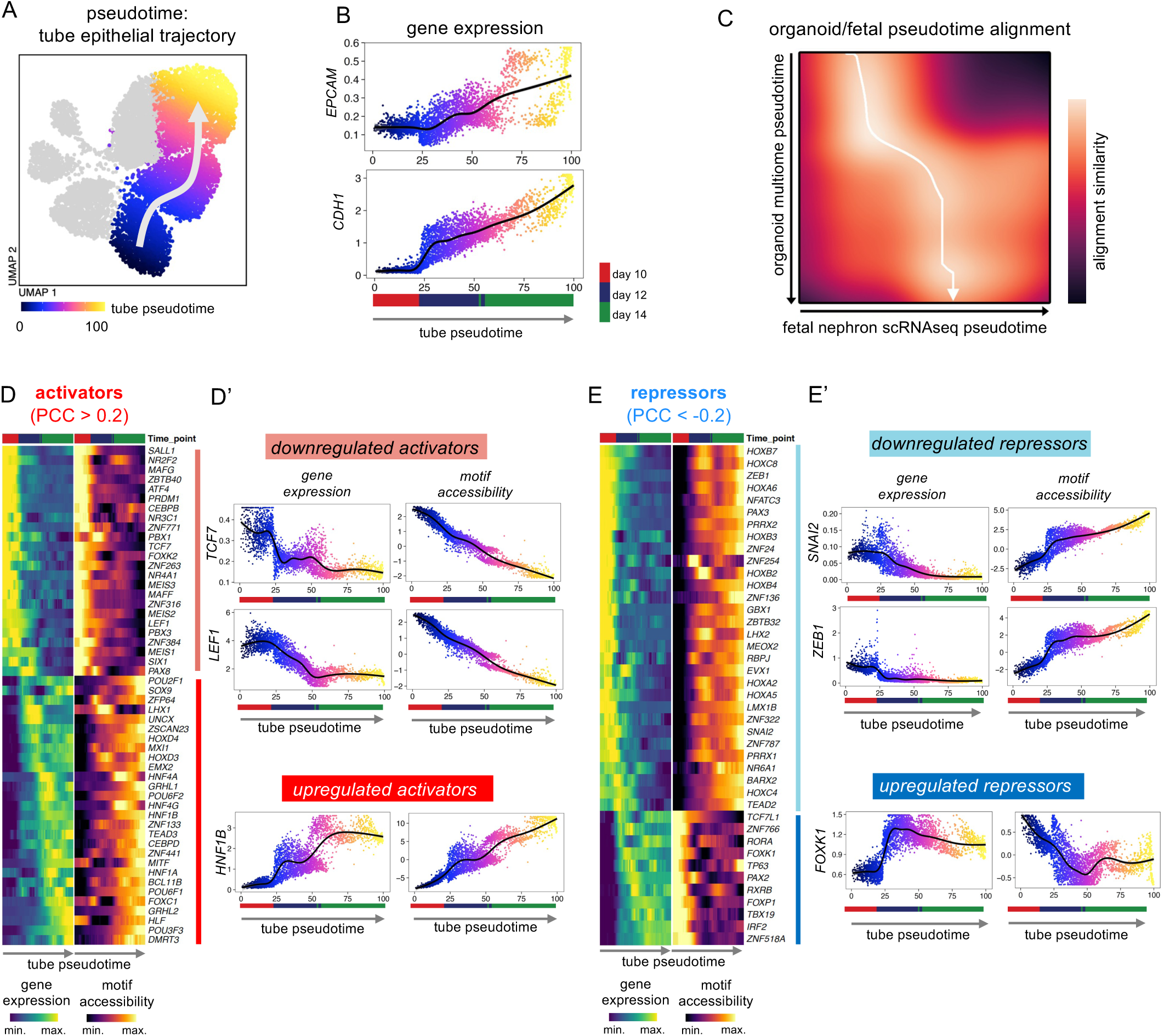
Dynamic activity of transcriptional activators and repressors drives renal MET. (A) UMAP plot of 9,147 multi-ome cells from batch 3 coloured according to tube epithelial pseudotime, as computed in ArchR (Granja et al., 2021). (B) Cells from batch 3 plotted according to tube epithelial pseudotime as in (A), and gene expression of *EPCAM* and *E-Cadherin/CDH1,* with time points indicated below. (C) Alignment of organoid tube epithelial pseudotime with fetal nephron pseudotime. The arrow indicates the path through the minima of the cost matrix where the trajectories best align. (D) Heat maps (left) of gene expression and corresponding motif accessibility along tube epithelial pseudotime for transcriptional activators (PCC gene expression vs motif accessibility along pseudotime > 0.2). Plots (right) of gene expression and motif accessibility of example downregulated activators *TCF7* and *LEF1*, and an example upregulated activator *HNF1B,* along tube epithelial pseudotime. (E) Heat maps (left) of gene expression and corresponding motif accessibility along tube epithelial pseudotime for transcriptional repressors (PCC gene expression vs motif accessibility along pseudotime < -0.2). Plots (right) of gene expression and motif accessibility of example downregulated repressors *SNAI2* and *ZEB1*, and an example upregulated repressor *FOXK1,* along tube epithelial pseudotime. See also Supplemental Fig. S6.

Two classes of activators were observed: the first showed downregulated gene expression during MET, which included *TCF7* and *LEF1*, consistent with attenuation of Wnt/β-Catenin signalling throughout MET (Fig. 6D). Interestingly, *LEF1* expression and motif accessibility were decreased at day 12 compared to day 10 despite continued presence of 1µM CHIR in the organoid protocol up to and including day 12 (Fig. 6D), suggesting a mechanism to limit the activity of Wnt/β-Catenin signalling downstream of GSK3. The second class of activators showed upregulated gene expression during MET, which included *HNF1B*, consistent with enrichment of HNF1A/B motifs in ATAC peaks at day 14 (Fig. 1J) and in agreement with the well-established role of HFN1B in renal development (Ferre & Igarashi, 2019).

Two classes of repressors were observed. One class showed decreased gene expression and increased motif accessibility during MET, which included the EMT-associated transcription factors (EMT-TFs) *SNAI2* and *ZEB1*. The second class of repressors showed upregulated gene expression with decreased motif accessibility during MET (Fig. 6E), which included the forkhead family transcription factor *FOXK1,* which was described as a transcriptional repressor in myoblasts and fibroblasts (Bowman et al., 2014). Corresponding analysis of batch 2 revealed similar trajectories for all these transcription factors during MET (Supplemental Fig. S6A-C).

Altogether, these analyses reveal that attenuation of Wnt/β-Catenin signalling during MET is associated with dynamic activity of specific activators and repressors that are likely to drive completion of human renal MET, and whose individual roles can now be probed using the organoid system.

### Attenuated Wnt/β-Catenin signalling permits completion of renal MET

Wnt/β-Catenin signalling specifies the nephron fate programme, a critical part of which, as we have shown, is PAX8-driven initiation of MET. However, it was reported that sustained Wnt/β-Catenin signalling blocks completion of renal MET (Park et al., 2012; Park et al., 2007), suggesting that Wnt/β-Catenin signalling inhibits the activity of transcription factors needed to properly organise epithelial polarity and junctions.

To identify transcription factors that drive completion of renal MET, we compared organoids cultured until day 14 in the normal protocol, with organoids in which completion of MET was blocked with sustained Wnt/β-Catenin signalling. As *LEF1* gene expression and motif accessibility were decreased at day 12 despite the presence of 1µM CHIR (Fig. 6E), we used 3µM CHIR to ensure sustained Wnt/β-Catenin signalling beyond day 12 (Fig. 7A).

**Figure 7.**
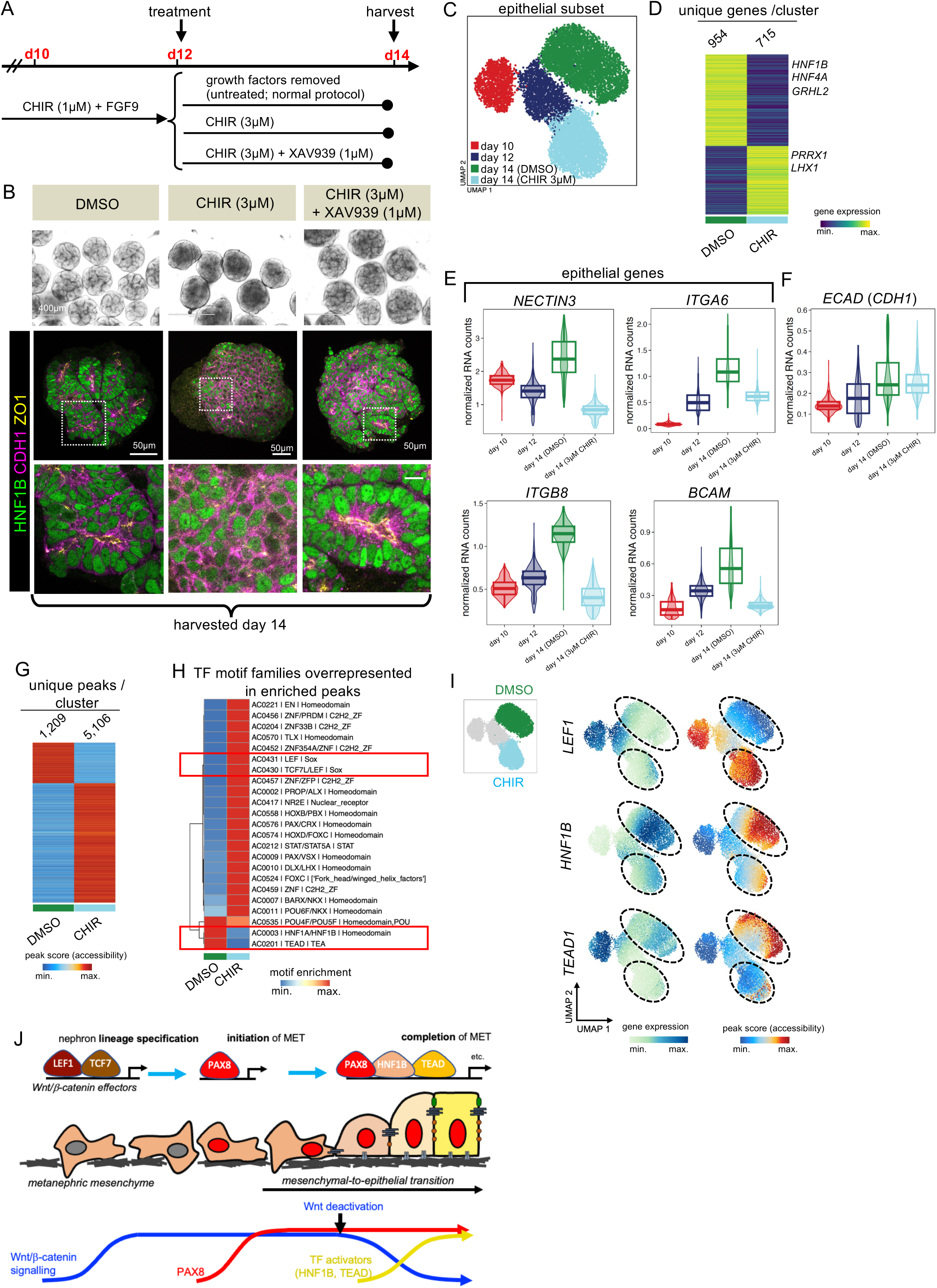
Attenuation of Wnt/β-Catenin signaling allows completion of MET via HNF1B and TEAD. (A) Schematic of experimental design to test the effect of persistent Wnt/β-Catenin pathway activation on completion of renal MET. (B) Organoids harvested at day 14 following treatments indicated in (A), with light microscopy images (top, scale bar 400µm), and immunofluorescence images showing expression of HNF1B (green), E-Cadherin/CDH1 (magenta) and ZO1 (yellow). Scale bars 50µm. (C) UMAP of cells from multi-ome batch 3 containing additional CHIR-treated group. (D) Heat map of marker genes expression levels for DMSO-treated or CHIR-treated epithelium at day 14 determined by snRNA-seq with representative markers annotated (log2 FC > 1, FDR < 0.01, two-sided Wilcoxon rank-sum test). (F, G) Gene expression violin plots of (E) *NECTIN3, ITGA6, ITGB8* and *BCAM,* and (F) *E-Cadherin/CDH1,* according to clusters annotated in (C). Box center line, median; limits, upper and lower quartiles; whiskers, 1.5x interquartile range. (G) Heat map of marker peaks of chromatin accessibility for DMSO-treated or CHIR-treated epithelium at day 14 determined by snATAC-seq (left, log2 FC > 1, FDR < 0.01, two-sided Wilcoxon rank-sum test). (H) Heat map of transcription factor motif archetypes (Vierstra et al., 2020) overrepresented in differentially accessible peaks from (G). Annotated are archetype codes, archetypal transcription factors and DNA-binding class. Highlighted in red are Wnt/β-Catenin transcriptional effectors LEF1 and TCF7/LEF, and HNF1A/HNF1B and TEAD family transcription factor motif archetypes. (I) UMAP plots highlighting DMSO-and CHIR-treated epithelium at day 14 (top), with UMAPs corresponding to gene expression (left) and motif accessibility (right) for *LEF1*, *HNF1B* and *TEAD*. (J) Our proposed model of the transcriptional control of morphogenetic events during human renal MET. See also Supplemental Figs S7 and S8.

Organoids at day 12 (when MET has initiated and early structures with apical-basal polarity are present, but adherens and tight junctions remain disorganised; Fig. 1B, C and Supplemental Figure S7A) were split into three groups. The control group received DMSO, one group received 3µM CHIR and one group received 3µM CHIR plus the tankyrase inhibitor XAV939 (1µM) to counteract the effects of CHIR and rescue any potential phenotype (Huang et al., 2009). Organoids were then cultured until day 14 (Fig. 7A). Control organoids at day 14 contained tubular epithelium that expressed HNF1B, with polarised localisation of E-Cadherin and ZO1 (Fig. 7B, Supplemental Fig. S7B). Strikingly, however, organoids in which Wnt/β-Catenin signalling was sustained lacked tubular structures, instead showing diffuse E-Cadherin and reduced HNF1B staining. Moreover, ZO1 staining was reduced, and where present was observed in sparse foci, indicating lack of proper organization of epithelial junctions (Fig. 7B, Supplemental Fig. S7B), similar to organoids at day 12 (Supplemental Fig. S7A). Addition of XAV939 rescued tubular morphology and resulted in organoids indistinguishable from control organoids (Fig. 7B, Supplemental Fig. S7B). Therefore, sustained Wnt/β-Catenin signalling blocked completion of renal MET in organoids, and prevented proper organisation of adherens and tight junctions. Notably, this effect was reversible, as a period of CHIR washout from day 14 to 17 produced organoids that contained tubular epithelial structures positive for HNF1B with apical-lateral localisation of E-Cadherin and ZO1, similar to control organoids at day 17 (Supplemental Fig. S7C and D). Thus, transcriptional mechanisms that drive proper epithelial junctional organisation are activated upon attenuation of Wnt/β-Catenin signalling.

To investigate transcription factors activated following Wnt/β-Catenin attenuation during MET, organoids treated from day 12 to 14 with 3µM were analysed by multi-ome sequencing in batch 3 (Supplemental Fig. S7E, Fig. 1H; see STAR METHODS) which together with the rest of batch 3 resulted in 13,502 cells that were visualised as a UMAP according to ATAC data (Supplemental Fig. S7F). The epithelial lineage (*PAX8^+^*, *EPCAM*^+^, *TWIST1^-^*, Supplemental Fig. S7G) was re-clustered, resulting in 4 clusters corresponding to day 10, day 12, and day 14 treated with DMSO or 3µM CHIR from day 12-14 (Fig. 7C).

Compared to control epithelial cells at day 14, treatment with 3µM CHIR from day 12-14 led to downregulation of genes involved in cell adhesion molecule binding and of transmembrane protein kinase and ion transport activity according to gene ontology analysis, suggesting defects in epithelial polarisation and segment differentiation (Fig. 7D, Supplemental Fig. 7H, Supplemental Table 6, Supplemental Table 7). CHIR treatment caused decreased expression of the cell adhesion molecule *NECTIN3,* which may contribute to establishment of epithelial polarity during normal kidney development (Brakeman et al., 2009), and to decreased expression of *ITGA6*, *ITGB8* and *BCAM* which are basolaterally-localised adhesion molecules; decreased expression of these genes may therefore contribute to disrupted epithelial polarity (Fig. 7E). Notably, gene expression for *E-Cadherin/CDH1* was unchanged (Fig. 7F). Therefore, sustained Wnt/β-Catenin signalling did not directly repress E-Cadherin expression but instead may dysregulate other adhesion molecules causing impaired establishment and maintenance of proper epithelial polarisation.

CHIR treatment led to differentially accessible chromatin regions in the epithelium compared to control at day 14 (Fig. 7G), which were then analysed for transcription factor motif archetypes to gain insights into transcription factors driving changes in epithelial morphology (Vierstra et al., 2020). This revealed enrichment of TCF7/LEF1 motifs in the CHIR-treated epithelium, confirming sustained Wnt/β-Catenin signalling (Fig. 7H). Notably, HNF1A/B and TEAD motif archetypes were selectively depleted following CHIR treatment (Fig. 7I). These data suggest that Wnt/β-Catenin signalling normally represses the activity of HNF1A/B and TEAD transcription factors, and so attenuation of Wnt/β-Catenin signalling permits their activity leading to completion of the MET programme via organisation and maintenance of epithelial polarity and junctions.

Interrogation of multi-ome data revealed loss of proximal and gain of distal tubule identity upon sustained Wnt/β-Catenin signalling (Supplemental Fig. S8A-D). Notably, CHIR treatment caused a complete loss of motif accessibility for the Notch signalling effector RBPJ (Supplemental Fig. S8D and E). Previous studies implicated Notch signalling in proximal fate differentiation during nephron segmentation (Duvall et al., 2022). Accordingly, CHIR treatment caused a reduction in gene expression of the Notch pathway ligands *NOTCH2* and *DLL1*, consistent with sustained Wnt/β-Catenin signalling inhibiting proximal differentiation via Notch inhibition (Supplemental Fig. S8F).

We conclude that attenuation of Wnt/β-Catenin signalling appears crucial to selectively allow HNF1A/B and TEAD family transcription factors to drive completion of MET, through their effects on the proper organisation and maintenance of epithelial junctions. However, the additional role of Wnt/β-Catenin signalling in proximal tube differentiation means its attenuation must be carefully balanced for normal nephrogenesis to proceed.

## Discussion

How the genome encodes form and hence function is still a major unresolved question in biology. Using human iPSC-derived renal organoids as a human developmental model, we comprehensively map the dynamics of transcription factor activity and resultant gene expression programmes at single cell resolution during the earliest acquisition of form and shape in human nephrogenesis, the renal MET. Our study elucidates the genetic cascade leading from fate commitment to morphogenesis and reveals that PAX8 directly activates a cell adhesion programme to initiate MET, and thus is a critical component of the Wnt/β-Catenin-dependent nephron fate programme in human kidney development. Subsequently, and following attenuation of Wnt/β-Catenin signalling, another tier of transcription factors is activated downstream of PAX8 that includes HNF1B and TEAD and is required to complete MET (Fig. 7J).

Pairing of transcriptome and chromatin accessibility profiling enables additional insights compared to use of either approach alone, one powerful example being the derivation of correlations between transcription factor gene expression and genome-wide accessibility of corresponding motifs. Using this approach, we identified numerous transcriptional activators and repressors potentially driving MET. Of note, we observed downregulation of the EMT-TFs SNAI2 (Slug), in agreement with previous studies in mouse (Boutet et al., 2006), and also of ZEB1; both SNAI2 and ZEB1 can directly bind to and repress epithelial genes including *E-Cadherin/CDH1* (Bolos et al., 2003; Eger et al., 2005; Ng-Blichfeldt & Röper, 2021), and therefore their downregulation may de-repress epithelial genes to allow progression and completion of MET. Our comprehensive multi-ome dataset identified several additional activators and repressors that may have a key role in MET, opening up many avenues for future work.

Our study is the first to show that PAX8 directly drives MET during human kidney development. To date, investigation into the precise role of PAX8 in renal MET has been hampered by partial redundancy with PAX2 *in vivo* (Bouchard et al., 2002; Narlis et al., 2007). Moreover, PAX8 is expressed *in vivo* not only in committed cap mesenchyme, but also in collecting duct and in earlier transient pro-and meso-nephric structures during renal development (Narlis et al., 2007). Therefore, while *PAX8* variants are associated with kidney defects in humans (albeit infrequently; (Carvalho et al., 2013; Meeus et al., 2004)), this could reflect a role in these other renal structures. The renal organoid system we used produces no ureteric bud derivatives, and appears to bypass uncommitted cap mesenchyme to instead recapitulate committed cap mesenchyme and subsequent MET where PAX8 is expressed *in vivo*, and so provides the unique opportunity to study the precise role of PAX8 in renal MET. Furthermore, our multi-ome approach enables unbiased identification of transcription factor regulons. With this approach we identified genes potentially regulated by PAX8, which by GO analysis were enriched for cell adhesion genes. In particular, activation of the key adhesion genes *CDH6* and *E-Cadherin/CDH1* by PAX8 helps to explain how PAX8 drives polarisation and initiates MET. The survey of PAX8-regulated genes presented here is likely to be of interest given the emerging evidence of PAX8-dependency in the development of adult human cancers including renal cell clear cell carcinoma and ovarian cancer (Bleu et al., 2021; Patel et al., 2022). All peak to gene links identified in our study are provided in Supplementary Table 8 as a resource to further explore gene programmes controlled by transcription factors driving human renal MET.

Our functional data diverge from previous studies where mice with germline *Pax8* null mutations had no apparent kidney phenotype, attributed to compensation by Pax2 *in vivo* (Bouchard et al., 2002; Narlis et al., 2007). PAX8 and PAX2, together with PAX5, comprise a PAX family subgroup and share a conserved DNA-binding paired domain, a partial homeodomain and an octapeptide domain, and have partially redundant functions *in vivo* (Bouchard et al., 2000). Our observation that *PAX2* is expressed in uncommitted cap mesenchyme in human fetal kidney *in vivo,* and persists upon commitment and subsequent MET, explains how PAX2 may compensate *in vivo* for loss of PAX8. Nonetheless, the presence of PAX2 in uncommitted cap mesenchyme with no apparent change at the onset of MET argues that PAX2 is unlikely to be directly driving renal MET as previously suggested (Rothenpieler & Dressler, 1993; Torres et al., 1995). PAX2 can form a complex with HOX paralogs and EYA1 to directly drive *Six2* expression, implicating PAX2 in the control of nephron progenitor self-renewal as opposed to commitment (Gong et al., 2007). Moreover, selective depletion of *Pax2* in *Six2*+ cells in mice caused ectopic transdifferentiation into renal stroma (Naiman et al., 2017), suggesting Pax2 may function to repress an interstitial fate within uncommitted cap mesenchyme. We speculate that specific co-factors are expressed upon commitment of cap mesenchyme together with PAX8 that allow PAX8 to activate the MET programme, whilst permitting these potentially divergent functions of PAX2. The multi-ome data presented herein provides a resource to aid future studies aimed at elucidating co-factors recruited by PAX8 and PAX2 to mediate gene regulation during renal MET.

It was known from studies by Park *et al*. that Wnt/β-Catenin signalling, though crucial to specify nephron fate, must act transiently for successful MET (Park et al., 2012; Park et al., 2007), but the reason underlying this was unknown. Applying multi-ome profiling to the renal organoid system allowed us to address this question, by revealing global transcription factor dynamics during MET as Wnt/β-Catenin signalling is attenuated, and enabling investigation into the impact of sustained Wnt/β-Catenin signalling with CHIR. Wnt/β-Catenin signalling has a close relationship with the epithelial state in part owing to the dual role of β-Catenin at adherens junctions, and as a transcriptional co-activator for TCF/LEF (Heuberger & Birchmeier, 2010). One possibility is that Wnt/β-Catenin activation sequesters β-Catenin in the nucleus, preventing recruitment to adherens junctions and thus inhibiting their proper organisation. However, our finding that sustained Wnt/β-Catenin signalling caused downregulation of genes involved in cell adhesion (e.g. *NECTIN3, ITGA6, ITGB8* and *BCAM*) argues that the Wnt-dependent transcriptional programme is broadly antagonistic to renal MET. Another possibility is that Wnt/β-Catenin signalling represses epithelial genes indirectly via EMT-TFs. Wnt/β-Catenin signalling can directly activate expression of EMT-TFs (Howe et al., 2003; Vallin et al., 2001) and also augment EMT-TF activity through post-translational stabilisation (Yook et al., 2005; Yook et al., 2006). Thus, attenuation of Wnt/β-Catenin signalling may lead to the *SNAI2* and *ZEB1* downregulation that we observed as renal MET proceeds, and their downregulation in turn could aid the completion of MET.

Our finding that Wnt/β-Catenin signalling becomes attenuated despite the continued presence of CHIR in the culture media until day 12 suggests an intrinsic mechanism to limit Wnt/β-Catenin signalling during MET. We found that the Hippo-YAP/TAZ pathway effector TEAD (Zhao et al., 2008) may mediate completion of renal MET, raising the intriguing possibility that biomechanical feedback could inhibit Wnt signalling. Increased tension at epithelial adherens junctions can sequester the serine/threonine kinases Lats1/2, thereby promoting YAP/TAZ accumulation and signalling (Dutta et al., 2018; Ibar et al., 2018; Rauskolb et al., 2014). In turn, YAP/TAZ can directly promote β-Catenin degradation to inhibit Wnt/β-Catenin signalling (Azzolin et al., 2014), together suggesting that Wnt signalling could be attenuated in response to tension establishment at maturing junctions, as would occur during renal MET. YAP/TEAD signalling itself must be carefully controlled for proper nephrogenesis, as deletion of *Lats1/2* in Six2^+^ cap mesenchyme cells in mice led to loss of nephrons, with mutant cells instead expressing mesenchymal markers (McNeill & Reginensi, 2017). Mechanisms leading to TEAD activation during renal MET remain to be determined, but one possibility could involve direct interactions between TEAD and PAX8, as has been observed in Müllerian epithelial cells (Elias et al., 2016). Another possibility suggested by our study could involve interaction between TEAD and HNF1B, which could co-operate to drive completion of renal MET. Accordingly, studies of mutant mice with kidney-specific deletion of HNF1B showed that early steps of renal MET are unaffected by HNF1B loss, with defects instead observed at comma-and S-shaped body stages, consistent with a role for HNF1B in the completion but not initiation of MET (Massa et al., 2013). HNF1B has also been reported to antagonise Wnt/β-Catenin signalling by both directly repressing *LEF1* gene expression and by competitively binding and antagonising TCF/LEF1-target genes (Chan et al., 2020; Chan et al., 2019). Increased expression of HNF1B, for example by direct activation by PAX8 as we observed, could thus provide another possible feedback mechanism to limit Wnt/β-Catenin signalling during renal MET.

Our study elucidates how the developing human kidney balances fate-commitment and morphogenesis, and offers a rich resource for the community to explore mechanisms controlling early nephron development in greater depth. Our study is therefore likely to have implications for our understanding of developmental kidney disease, but also aberrant epithelial plasticity incurred by acute injury to renal tubular epithelium in adulthood. Acute kidney injury is thought to re-engage developmental processes, particularly activation of Wnt/β-Catenin signalling leading to partial EMT, cellular responses which contribute to tubular repair but can also pre-dispose to chronic kidney disease (Schunk et al., 2021). A more complete picture of early events in renal morphogenesis as presented here is therefore likely to aid a better understanding of how to modulate the response to acute kidney injury for successful kidney regenerative strategies.

## Acknowledgments

The authors would like to thank the following people: members of the lab and Cell Biology division for input on the manuscript; the Genomics Facility at Cancer Research UK, Cambridge, UK. K.R., J.P.N.B. were supported by the Medical Research Council (file reference number U105178780). M.R.C. and B.J.S. were supported by an NIHR Research Professorship (RP-2017-08-ST2-002), B.J.S by a Wellcome Trust Clinical Training Fellowship (216366/Z/19/Z), and M.R.C. by a Wellcome Trust Investigator Award (220268/Z/20/Z) and Chan Zuckerberg Foundation Kidney Seed Network Award. The Clatworthy lab utilises infrastructure supported by the National Institute of Health Research (NIHR) Cambridge Biomedical Research Centre (NIHR203312). The views expressed are those of the authors and not necessarily those of the NIHR or the Department of Health and Social Care. This work was supported by the Medical Research Council, as part of United Kingdom Research and Innovation (also known as UK Research and Innovation) [MRC file reference number U105178780]. For the purpose of open access, the author has applied a CC BY public copyright licence to any Author Accepted Manuscript version arising. This project is supported through a research collaboration between AstraZeneca UK Limited and the Medical Research Council.

## Author contribution

Conceptualisation, K.R., J.M.W., J.P.N.B.; Methodology, J.P.N.B., B.J.S., M.R.C..; Investigation, J.P.N.B., B.J.S..; Writing-Original Draft, K.R., J.P.N.B.; Funding Acquisition, K.R., J.M.W, M.R.C.

## Materials and Methods

### Key resources table

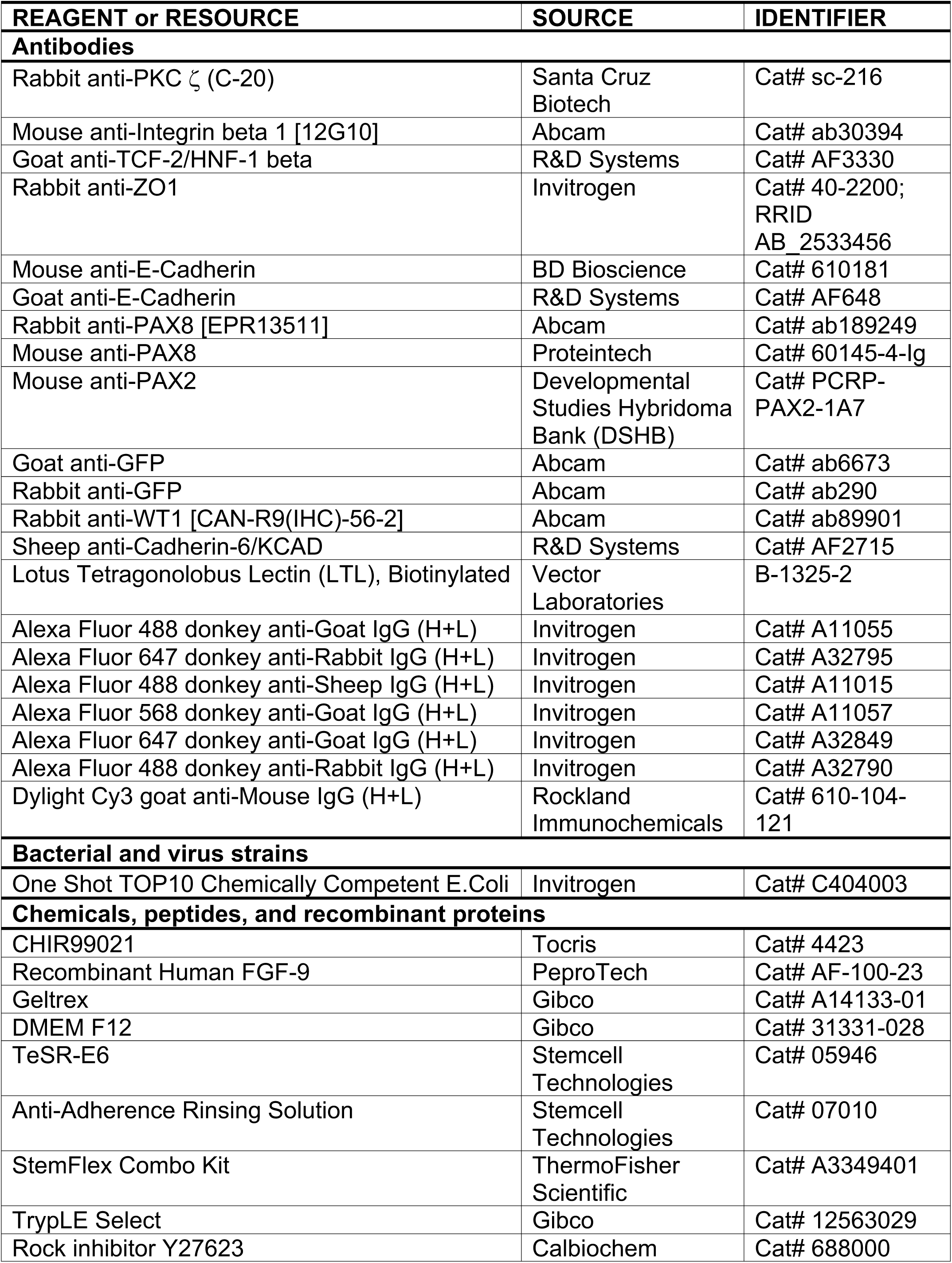
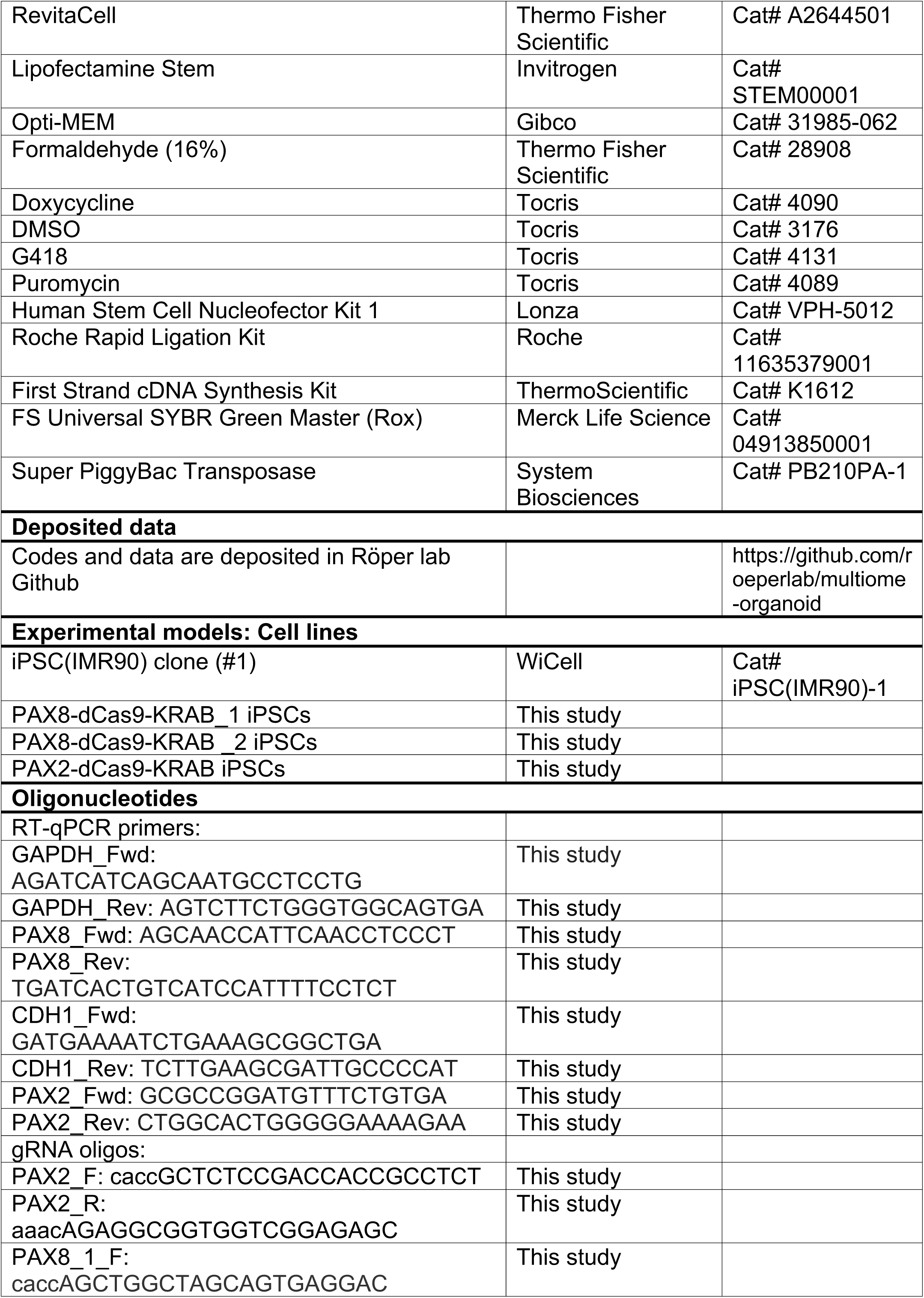
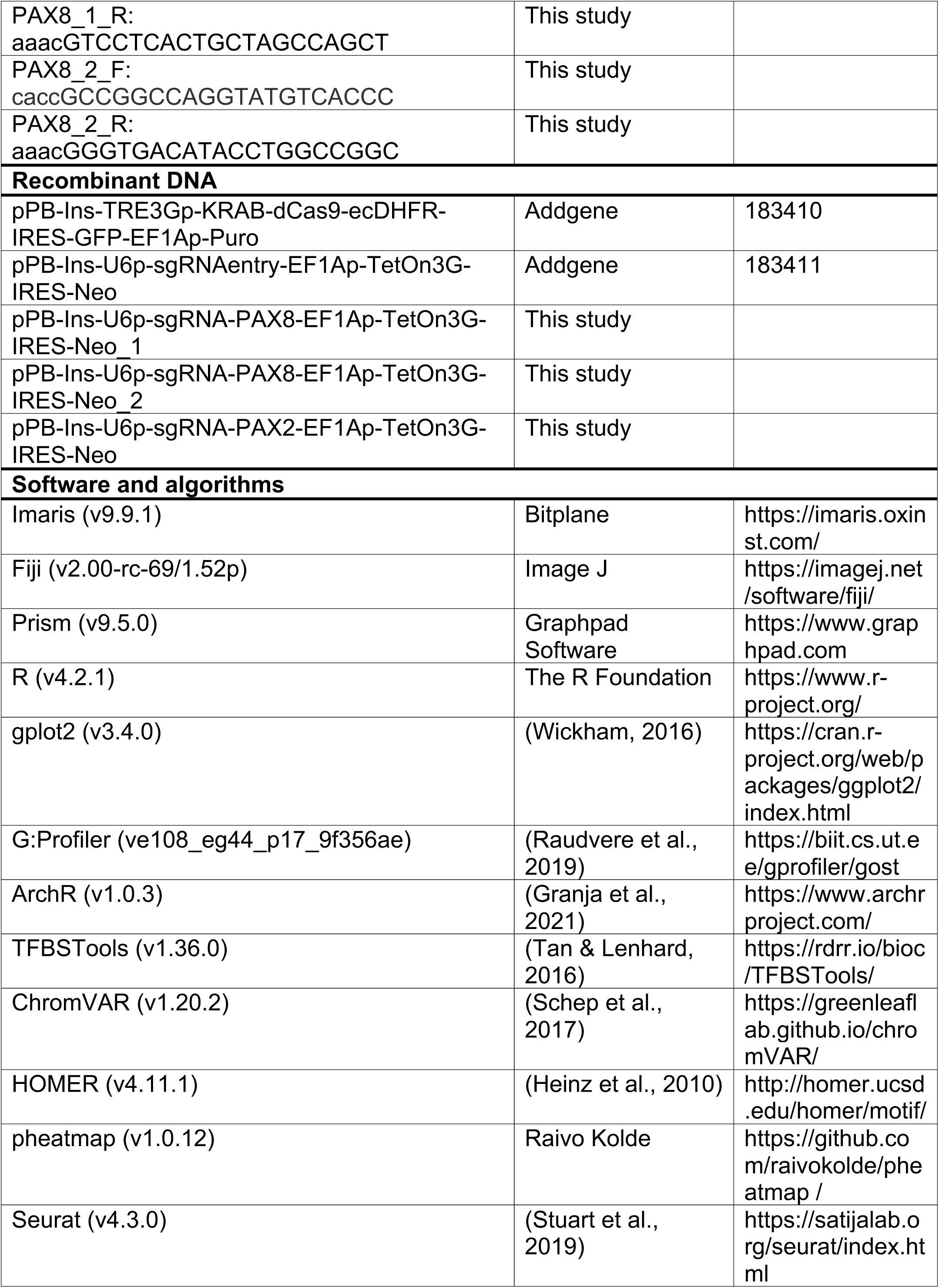
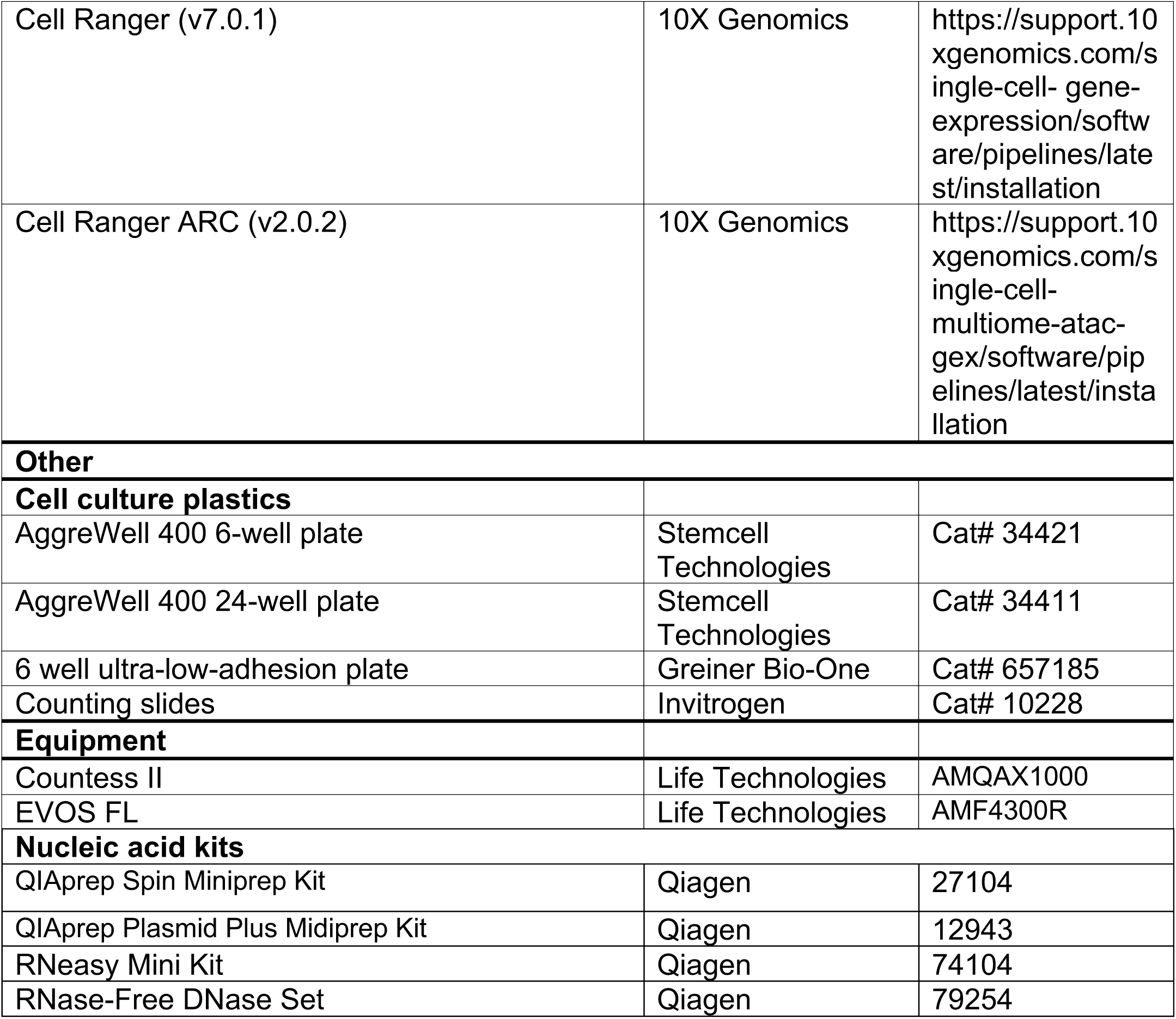

### Materials Availability Statement

All unique/stable reagents generated in this study are available from the lead contact with a completed materials transfer agreement.

### Contact for Reagent and Resource Sharing

Katja Röper

## EXPERIMENTAL METHOD AND SUBJECT DETAILS

### Cell lines

iPS(IMR90)-1 (IMR-90-1) cells were obtained from WiCell, and were maintained in StemFlex (ThermoFisher, A3349401) containing 1x Antibiotic-Antimycotic (Anti-/Anti-; Gibco, 15240062) on Matrigel-or Geltrex-coated plates. Cells were passaged every 3-4 days using EDTA and re-seeded in StemFlex containing 1x Anti-/Anti-and 10µM ROCK inhibitor Y27623 (ROCKi; Calbiochem, 688000). PAX8-dCas9-KRAB-puro (PAX8-dCas9-KRAB_1 and PAX8-dCas9-KRAB_2) iPSCs and PAX2-dCas9-KRAB-puro (PAX2-dCas9-KRAB) iPSCs, generated as described below, were maintained in StemFlex containing 1x Anti-/Anti-, 0.5 µg/mL puromycin (Tocris, 4089) and 200 µg/mL G418 (Tocris, 4131). Cells were passaged every 3-4 days using EDTA and re-seeded in StemFlex containing 1x Anti-/Anti-, 10µM ROCKi, 0.5 µg/mL puromycin (Tocris, 4089) and 200 µg/mL G418 (Tocris, 4131). All cell cultures were incubated at 37°C with 5% CO_2_.

## METHOD DETAILS

### Plasmid constructs

pPB-Ins-U6p-sgRNAentry-EF1Ap-TetOn3G-IRES-Neo was a gift from Azim Surani (Addgene plasmid # 183411; http://n2t.net/addgene:183411 ; RRID:Addgene_183411). pPB-Ins-TRE3Gp-KRAB-dCas9-ecDHFR-IRES-GFP-EF1Ap-Puro was a gift from Azim Surani (Addgene plasmid # 183410 ; http://n2t.net/addgene:183410 ; RRID:Addgene_183410). All oligos used for cloning were ordered using the custom DNA oligo synthesis service from Merck and are listed in the “Oligonucleotides” section of the Key resources table. Specific gRNA sequences for dCas9-KRAB mediated gene repression were designed by identifying putative transcription start sites (TSSs) for target genes according to FANTOM CAGE data publicly available on ZENBU (Severin et al., 2014), then the Wellcome Sanger Institute Genome Editing (WGE) CRISPR Finder (Hodgkins et al., 2015) was used to identify CRISPR sequences 50-100bp downstream of a putative TSS for optimal gene knockdown as previously described (Gilbert et al., 2014). At least 2 gRNA sequences were tested per TSS. For generation of gRNA expression vectors, complementary gRNA oligos were annealed and ligated into pPB-Ins-U6p-sgRNAentry-EF1Ap-TetOn3G-IRES-Neo (Addgene plasmid, 183411) by Esp3I digestion (NEB, R0734) followed by ligation with Rapid DNA Ligation Kit (Roche, 11635379001). TOP10 chemically competent E. coli (Thermo Fisher, C404010) were used for transformation. Successful transformants were purified using QIAprep Spin Miniprep Kit (Qiagen, 27104) or QIAprep Plasmid Plus Midiprep Kit (Qiagen, 12943), then validated with Sanger sequencing.

### Quantitative reverse transcriptase PCR (RT-qPCR)

To isolate RNA for RT-qPCR analysis, cells/organoids were washed twice with PBS and then lysed in 350 µl RLT lysis buffer prior to RNA isolation with RNeasy Mini Kit (Qiagen, 74104) with on-column Dnase digestion using the Rnase-free Dnase set (Qiagen, 79254). Complementary DNA (cDNA) was generated using First Strand cDNA Synthesis Kit (Thermo Scientific, K1612) with 1 µg RNA for each sample. RT-qPCR was performed using SYBR Green (Roche, FSUSGMMRO) with forward and reverse primers diluted to final concentration of 1µM per reaction, with 5.2 µl diluted cDNA, in MicroAmp Optical 384-well Reaction Plate with Barcode (Applied Biosystems, 43309849) with MicroAmp Optical Adhesive Film (Applied Biosystems, 4311971) overlaid. Reactions were performed in an Applied Biosystems ViiA7 machine with the following cycling conditions: Hold for 10 minutes at 95°C, then 10 seconds at 95°C and 20 seconds at 58°C repeated for 40 cycles, followed by a melt curve stage. The DDCt method (Livak & Schmittgen, 2001) was used to analyse PCR data using *GAPDH* as a housekeeping gene. RT-qPCR data were analysed using Prism (Graphpad, v9.5.0).

### Transgenic cell lines

For establishment of PAX8-dCas9-KRAB_1, PAX8-dCas9-KRAB_2 and PAX2-dCas9-KRAB lines, 5×10^5^ IMR-90-1 cells were seeded into Matrigel-or Geltrex-coated 6-well plates in StemFlex with ROCKi, and the following day were washed with PBS and incubated in E8 media (Gibco, A1517001) for at least 1 hour. For each well of a 6-well plate, cells were then transfected with the plasmids Super piggyBac Transposase expression vector (Stratech, PB210PA-1-SBI) (500ng), pPB-Ins-U6p-sgRNA-PAX8-EF1Ap-TetOn3G-IRES-Neo_1, pPB-Ins-U6p-sgRNA-PAX8-EF1Ap-TetOn3G-IRES-Neo_2, or pPB-Ins-U6p-sgRNA-PAX2-EF1Ap-TetOn3G-IRES-Neo gRNA expression plasmids (1µg) and pPB-Ins-TRE3Gp-KRAB-dCas9-ecDHFR-IRES-GFP-EF1Ap-Puro (1.5µg) mixed with 10µl Lipofectamine Stem transfection reagent (Life Technologies, STEM00003) in 500µl Opti-MEM with 1x Revitacell supplement for 20 minutes followed by addition of StemFlex with 1x Revitacell supplement. The following day, media were replaced with fresh StemFlex with 1x Revitacell supplement. 48 hours later, media were refreshed with fresh StemFlex with 1x Revitacell supplement and containing antibiotics 0.5 µg/mL puromycin and 200 µg/mL G418 for selection. Media were refreshed with selection media every 2-3 days thereafter. Around a week later, surviving colonies for each well were dissociated together (effectively pooling multiple colonies) and re-seeded into 2 wells. Once cells reached ∼80% confluence, knockdown validation. For knockdown validation, transgenic lines were seeded for organoid generation into 3 wells, differentiated to Day 7 with 1 well additionally treated with 1µM Dox, and another well treated with 4µM Dox, then all harvested for RNA extraction. For knockdown validation, RT-qPCR was performed with primers specific to the gene of interest according to the “Quantitative reverse transcriptase PCR (qRT-PCR)” section below. PAX8-dCas9-KRAB_1 and PAX8-dCas9-KRAB_2 iPSCs showed similar knockdown efficiency and knockdown phentoypes, so only data from PAX8-dCas9-KRAB_1 are presented in this paper.

Piggybac transposase facilitates the integration of a transposon-containing donor cassette specifically at ’TTAA’ sites which are randomly dispersed in the genome, and therefore the activity of the Dox-inducible dCas9 cassette may be subject to genomic position effects leading to heterogenous responsiveness to Dox and consequently variable knockdown efficiency. To improve knockdown efficiency of PAX8-dCas9-KRAB_1 and PAX2-dCas9-KRAB iPSCs, cells were incubated with 1µM Dox for 48 hours to induce the expression of the dCas9-KRAB cassette which includes cytoplasmic GFP as a readout for Dox responsiveness. Cells were then dissociated to single cells, and the top 20% of GFP^+^ cells isolated by flow cytometry, followed by culture on Matrigel-or Geltrex-coated plates in StemFlex containing 1x Anti-/Anti-and 1x Revitacell (Gibco, A2644501). Top 20% sorted PAX2-dCas9-KRAB cells were used to generate all PAX2-dCas9-KRAB knockdown data presented in this paper (Supplemental Fig. S5D and E). Top 20% sorted PAX8-dCas9-KRAB_1 cells differentiated to organoids showed near complete knockdown of *PAX8* and completely lacked epithelial structures (data not shown).

### Kidney organoid generation

Kidney organoids were generated using a previously published protocol with modifications (Kumar et al., 2019). Human iPSCs were grown to ∼70% confluency and dissociated with EDTA (Day -1), counted using Countess II (Life Technologies, AMQA×1000), then seeded evenly onto Geltrex-coated 6-well plates at 100,000 cells per cm^2^ in 2ml StemFlex with ROCKi. The following day (Day 0), cells were washed with PBS, then incubated in 2ml fresh organoid induction media A: TeSR-E6 (Stemcell Technologies, 05946) with 1x Anti-/Anti-and 6µM CHIR99021 (Tocris, 4423). Media were refreshed on Day 3. On Day 4, media were replaced with organoid induction media B: TeSR-E6 with 1x Anti-/Anti-, 1µM CHIR and 200ng/ml FGF9 (PeproTech, AF-100-23). Media were refreshed on Day 6. On Day 7, cells were washed with PBS and dissociated with TrypLE (3 mins at 37°C), spun down and resuspended in organoid induction media B with 10µM ROCKi, and counted. Cells were then seeded into Aggrewell-400 24-well plates (Stemcell Technologies, 34411) or 6-well plates (Stemcell Technologies, 34421) that were pre-coated with Anti-adherence solution (Stemcell Technologies, 07010), at 1.2×10^6^ cells per well (24-well) or 7×10^6^ per well (6-well). Plates were centrifuged at 200g for 5 mins, then incubated at 37°C with 5% CO2. At Day 10, the resulting organoids were gently washed out of Aggrewell plates, washed once with PBS, and transferred at ∼3000 organoids per well into 6-well ultra-low-adhesion plates (Greiner Bio-One, 657185) in fresh organoid induction media B, and incubated on a shaker at 100rpm, at 37°C with 5% CO2. At Day 12, media were replaced with TeSR-E6 with 1x Anti-/Anti-. Media were refreshed with TeSR-E6 with 1x Anti-/Anti-at Day 14, and every 2-3 days thereafter.

This protocol first generates intermediate mesoderm as indicated by the induction of *OSR1* gene expression (peaking at Day 3), followed by generation of nephron progenitor cells, indicated by the induction of *SALL1* gene expression (peaking at Day 5; Supplementary Fig. 1A).

### Immunofluorescence analysis

To harvest for immunofluorescence (IF) analysis, organoids were washed once with PBS, fixed in 3% PFA overnight at 4°C, then washed in PBS (3×10 min) and stored in PBS with 0.01% sodium azide. Prior to antibody staining, samples were incubated in blocking buffer overnight at 4°C (PBS containing 0.25% Triton-X and 5% BSA). Samples were incubated in primary antibodies diluted in wash buffer (PBS containing 0.1% Triton-X and 5% BSA) at the indicated concentrations overnight at 4°C. Next, samples were washed in wash buffer (3×15 min at room temperature) and incubated in secondary antibodies diluted in wash buffer at the indicated concentrations, with DAPI, overnight at 4°C. Then, samples were washed in wash buffer (3×15 min at room temperature) and mounted on glass slides in Vectorshield (Vector Labs, H1000) in 120µm spacers (Invitrogen, S24735) with coverslips overlaid.

### Imaging and image analysis

Brightfield images were taken using an EVOS FL microscope (Life Technologies, AMF4300R). Confocal images were taken using a Zeiss LSM 710 or 780, or a Nikon W1 Spinning Disk microscope. Image files were analysed using Fiji (Image J) or Imaris (Bitplane, Oxford Instruments, v9.9.1). To quantify E-Cadherin-positive volume, confocal z-stacks of individual organoids were first obtained using a Nikon W1 Spinning Disk microscope with imaging at 1 µm intervals, the image files were then imported into Imaris, and the surfaces option was used to volume render and quantify the E-Cadherin signal. All images presented are representative of at least 2 independent organoid batches. Unless otherwise indicated, immunofluorescence images shown are representative single confocal sections.

### Antibodies

Primary antibodies used for protein detection, with their corresponding dilutions for immunofluorescence (IF) were as follows: rabbit anti-PKC ζ (C-20) (Santa Cruz Biotech, sc-216, 1:200), mouse anti-Integrin beta 1 [12G10] (Abcam, ab30394, 1:200), goat anti-TCF-2/HNF-1 beta (R&D Systems, AF3330, 1:200), rabbit anti-ZO1 (Invitrogen, 40-2200, 1:200), mouse anti-E-Cadherin (BD Bioscience, 610181, 1:200), goat anti-E-Cadherin (R&D Systems, AF648, 1:200), rabbit anti-PAX8 [EPR13511] (Abcam, ab189249, 1:200), mouse anti-PAX8 (Proteintech, 60145-4-Ig, 1:200), mouse anti-PAX2 (Developmental Studies Hybridoma Bank (DSHB), PCRP-PAX2-1A7, 1:20), rabbit anti-GFP (Abcam, ab290, 1:200), rabbit anti-WT1 [CAN-R9(IHC)-56-2] (Abcam, ab89901, 1:200), sheep anti-Cadherin-6/KCAD (R&D Systems, AF2715, 1:200). Biotinylated Lotus Tetragonolobus Lectin (LTL; Vector Laboratories, B-1325-2, 1:200) was used to stain proximal tube cells. Alexafluor 488, 568 and 647 (Invitrogen), and Dylight Cy3 (Rockland Immunochemicals) secondary antibodies were used for detection of primary antibodies.

### RNA-seq sample preparation and sequencing

Organoids from batch 1 (days 10, 14, 24) and from batches 2 and 3 (day 24) were collected for single cell RNA-seq. For each sample, ∼5000 organoids were collected in 15ml tubes, washed once with PBS then dissociated to single cells in 500µL Bacillus Licheniformis protease (Sigma, P5380) on ice for 20 min (Adam et al., 2017), with gentle agitation by pipetting every 2 min, followed by resuspension in PBS with 10% FBS to neutralise the protease. Cells were centrifuged at 300g for 5 min, then resuspended in PBS with 10% FBS and filtered through a 30µm strainer prior to centrifugation and final resuspension in PBS with 0.5% BSA. Cells were counted with Countess II (Life Technologies) to ensure cell viability > 90% and to achieve a targeted cell recovery of 1,600 cells per sample for batches 1 and 2, and 5,000 cells per sample for batch 3. Cells were processed for library preparation using a 10X Chromium sc 3’mRNA kit (v3, 10X Genomics) at the Cancer Research UK Genomics Facility, Cambridge, UK.

### Multiome-seq sample preparation and sequencing

Organoids from batch 2 (days 10, 12, and 14) and batch 3 (days 10, 12, 14 DMSO, and 14 CHIR), were collected for single nucleus Multiome-seq. Single cell preparations were generated as above, then further processed for single nucleus isolation by centrifugation at 300g for 4 min at 12°C and resuspension in 100µL freshly prepared ice-cold lysis buffer (10mM Tris-HCL, 10mM NaCl, 3mM MgCl2, 0.1% Tween-20, 1 % BSA, 1mM DTT, 1 U/µL Protector RNase Inhibitor (Sigma, 3335402001), 0.1% Nonidet P40 Substitute (Sigma, 74385) and 0.01% Digitonin in nuclease-free H2O) followed by incubation on ice for 5 min. Lysis was halted by addition of 900µL chilled wash buffer (10mM Tris-HCL, 10mM NaCl, 3mM MgCl2, 0.1% Tween-20, 1 % BSA, 1mM DTT, 1 U/µL Protector RNase Inhibitor in H2O), followed by 3 rounds of centrifugation at 500g for 5 min at 4°C and resuspension in chilled wash buffer, with the final resuspension in 50µL chilled 1x Nuclei Buffer (10X Genomics, 2000153) with 1mM DTT and 1 U/µL Protector RNase Inhibitor in H2O, according to the 10x Genomics demonstrated protocol for Nuclei Isolation for Single Cell Multiome (ATAC + GEX) sequencing (CG000365, RevA). After resuspension in 1x Nuclei Buffer, nuclei were stained with Trypan Blue and counted using a Countess II (Life Technologies) to confirm complete lysis and to achieve a targeted nuclei recovery of 1,600 nuclei per sample for batch 2, and 5,000 nuclei per sample for batch 3. Nuclei were processed for library preparation using a 10X Chromium scMultiome kit (10X Genomics) at the Cancer Research UK Genomics Facility.

All samples from a given batch were sequenced together (RNA and ATAC libraries were sequenced separately) on an Illumina NovaSeq 6000 to an aimed depth of 100,000 – 150,000 reads per cell.

### Single cell RNA-seq and multi-ome (paired single nucleus RNA-seq and single nucleus ATAC-seq) data preprocessing

De-multiplexed scRNA-seq FASTQ files were run in the Cell Ranger pipeline (10X Genomics, v7.0.1) to produce barcoded count matrices of RNA data. Paired de-multiplexed snRNA-and snATAC-seq FASTQ files were run in the Cell Ranger ARC pipeline (10X Genomics, v2.0.2) to produce barcoded count matrices of RNA data and fragment files of ATAC data. All reads were aligned to GRCh38/hg38.

### scRNA-seq analysis

For analysis of organoid transcriptomes at days 10, 14 and 24 combined across 3 batches (Supplemental Fig. S1C-N), filtered feature-barcode matrices of scRNA-seq data (batch 1, days 10, 14, and 24; batches 2 and 3, day 24) and snRNA-seq data (batches 2 and 3, days 10 and 14) were converted into individual Seurat objects using the Seurat R package (version 4.3.0) (Stuart et al., 2019) and quality control filtered for cells containing > 1000 genes (> 2500 genes for batch 1, day 14) and < 15% mitochondrial reads. All samples were then merged into a single Seurat object, then split using the SplitObject function according to batch. Gene expression counts were normalized for each batch separately using the NormalizeData function. Features that were repeatedly variable across batches were identified using the SelectIntegrationFeatures function, which were then used to scale each batch separately using the ScaleData function and to run PCA. Samples within each batch were integrated using reciprocal PCA; the 3 integrated batches were then integrated using canonical correlation analysis. Graph-based clustering was performed on the final integrated object using the top 44 principal components at a resolution of 0.02 to generate 2 clusters.

Marker genes for each cluster were identified using the FindMarkers function with a minimum fraction of 0.2 and a minimum difference fraction of 0.1, which were used to annotate the clusters as either epithelial or stromal lineages. The epithelial lineage cluster was then subset and re-clustered using the top 28 principal components at a resolution 0.2. Marker genes for each cluster were identified using the “FindAllMarkers” function with a minimum fraction of 0.5 and a log2 fold change of 1. Cluster identities were then manually annotated by querying resulting marker genes using the Human Nephrogenesis Atlas (https://sckidney.flatironinstitute.org/) (Lindstrom et al., 2021) and by assessing cell cycle phase scored using the “CellCycleScoring” function in Seurat (Supplemental. Fig S1K)

### Integration of human fetal and organoid scRNA-seq data

Reference data from (Stewart et al., 2019) was pre-processed using the scanpy framework, and pseudotime calculated using scFates (Faure et al., 2023). Using reference PCA embeddings, organoid scRNAseq data was asymmetrically integrated using the “sc.tl.ingest” function. Label prediction between scFates milestones in fetal nephron data and scRNAseq profiles from organoids was performed using a multilayer perceptron in sklearn as previously described (Stewart et al., 2023). Pseudotime trajectories were aligned between fetal data and organoid data by first calculating genes that vary significantly over fetal nephron pseudotime with switchDE (Campbell & Yau, 2017), taking genes with a q value < 0.05.

Trajectories were aligned using cellAlign (Alpert et al., 2018), using 500 interpolated nodes.

### Multi-ome (paired snRNA-seq and snATAC-seq) data analysis

Paired snRNA-seq and snATAC-seq datasets (batches 2 and 3, days 10, 12, and 14) were analyzed using ArchR (version 1.0.3) (Corces et al., 2018; Granja et al., 2021) with default settings for all functions used unless otherwise stated. The RNA count matrices and ATAC fragment files output from Cell Ranger were used to generate Arrow files for each sample. Initially, all Arrow files (batches 2 and 3, all time points) were combined into one ArchR project, and batch correction with Harmony (Korsunsky et al., 2019) as implemented in ArchR was applied but was found to be inadequate to resolve batch effects without losing biological information, and therefore batches were analysed independently. Thus, for each batch separately, Arrow files for days 10, 12 and 14 were combined into an ArchR project and quality control filtered for nuclei with 1,000-10,000 genes, >5,000 RNA transcripts, TSS enrichment >6, and >2,500 ATAC fragments. Dimensionality reduction was performed with the “addIterativeLSI” method for ATAC and RNA data separately at a resolution of 0.2, using 2500 variable features for RNA data and 25000 variable features for ATAC data. For batch 3 only, Harmony (Korsunsky et al., 2019) was used to remove batch effects between samples but preserve the structure of the data with regards to time point (groupBy = “Sample”, theta = 120, lambda = 0.3). For each batch, UMAP dimensionality reduction was then performed (minDist = 0.8, dims = c(1:30)). Clustering was performed at a resolution of 0.9, and poor quality and stressed cell clusters were identified and removed from the dataset. The remaining cells were again subjected to UMAP dimensionality reduction (minDist = 0.8, dims = c(1:28)) followed by clustering at a resolution of 0.1 to generate 6 clusters corresponding to stromal and epithelial lineages each at day 10, 12 and 14 for batch 3; as we were focussing on the epithelial lineage, all stromal clusters were merged into 1 cluster for subsequent analysis (Figure 1E). Analogous processing produced 5 clusters in batch 2 data which were used for downstream analysis including pseudotime trajectory inference (Supplemental Fig. S4B and S6A-C). Gene expression values for visualization were imputed using MAGIC (van Dijk et al., 2018) and embedding plots were generated with the “plotEmbedding” function (quantCut = c(0.05,0.95)). Gene ontology analysis was performed by analysing gene lists in g:Profiler (https://biit.cs.ut.ee/gprofiler/gost) (Raudvere et al., 2019).

ATAC data were used to generate pseudobulk replicates with the “addGroupCoverages” and “addReproduciblePeakSet” functions. Chromatin accessibility peaks on chromosomes 1-22 and X and outside of blacklist regions were called with MACS2 (Zhang et al., 2008). To calculate enrichment of chromatin accessibility of transcription factor motifs on a per cell basis, ChromVAR (Schep et al., 2017) was run with “addDeviationsMatrix”, using motif sets from JASPAR 2020 core and unvalidated collections (as currently supported within ArchR) (Fornes et al., 2020) together with PAX2, MAFB, WT1 and TWIST2 motifs, which were not present in the JASPAR 2020 human collection and so were manually generated using position frequency matrices of mouse motifs from JASPAR 2020 and then converted into position weight matrices using PWMatrixList from the TFBSTools package (v1.36.0) (Tan & Lenhard, 2016). The SALL1 motif was taken from a previously published mouse Sall1 motif sequence (Yamashita et al., 2007). To visualize motif deviations, scores were imputed using MAGIC, and the “plotEmbedding” function was used (quantCut = c(0.05,0.95)). Marker genes for clusters as indicated in the text were identified using the “getMarkerFeatures” function using the Wilcoxon test and correcting for TSS Enrichment, and ATAC and RNA sequencing depth and filtered using the “getMarkers” function (cutOff =“FDR <= 0.01 & Log2FC >= 1”). Marker peaks of clusters as indicated in the text were identified using the “getMarkerFeatures” function using the Wilcoxon test and correcting for TSS Enrichment, and ATAC and RNA sequencing depth, followed by filtering with the “getMarkers” function (cutOff = “FDR <= 0.01 & Log2FC >= 1”). Enrichment of transcription factor motifs within marker peaks for each cluster was determined using the hypergeometric test using the “peakAnnoEnrichment” function based on position frequency matrices from the JASPAR 2020 motif set, with all peaks used as a background.

Vierstra transcription factor archetypes (Vierstra et al., 2020) were added using “addMotifAnnotations” (motifSet = “vierstra”). Once added, enrichment of Vierstra archetypes was determined using the “getMarkerFeatures” function using the Wilcoxon test and correcting for TSS Enrichment, and ATAC and RNA sequencing depth, followed by filtering with the “getMarkers” function (cutOff = “FDR <= 0.01 & Log2FC >= 1”). Transcription factor footprinting was performed and visualized using the ArchR functions “getFootprints” and “plotFootprints”, with the Tn5 insertion signal subtracted from footprinting signals prior to plotting to correct for Tn5 insertion bias (normMethod = “subtract”).

### Correlating transcription factor gene expression and corresponding motif accessibility

To identify activator and repressor transcription factors, correlations between transcription factor gene expression and chromatin accessibility of corresponding motifs were derived using the “correlateMatrices” function. Resulting transcription factors were filtered according to a correlation adjusted p value of 0.001, and further filtered using the “getMarkers” function by correlation value (>0.2 or < -0.2 to classify transcription factors as activators or repressors, respectively) and Log2fold change in the epithelial lineage compared to the stromal lineage (>1.5 or <-1.5 for epithelial enriched or stromal enriched transcription factors, respectively) (Figure 2E).

### Peak to gene links and regulon analysis

Peak to gene links were defined as chromatin accessibility peaks within 500kb upstream or downstream of a gene TSS with accessibility that correlates with expression of that gene, and were identified using the “addPeak2GeneLinks” function (maxDist = 500000) and filtered according to false discovery rate <1×10^-4^ and correlation value >0.4 (Supplemental Fig. S5F). Filtered peak to gene links were then analysed for overrepresentation of specific transcription factor motifs within HOMER (v4.11.1) (Heinz et al., 2010) using the “findMotifsGenome.pl” function with HOMER Motif files as input. Sequencing tracks of chromatin accessibility were generated in ArchR using the “plotBrowserTrack” function and were normalized by the total number of reads in TSS regions.

### Pseudotime trajectory analysis

To analyse dynamics of transcription factor activity along the renal mesenchymal-to-epithelial transition, pseudotime analysis was performed using ArchR. For batch 3 (figure 5), the ArchR object was clustered at a resolution of 0.3, and the “addTrajectory” function applied using the following user-defined trajectory as a guide for the supervised trajectory analysis: “Cluster 3”, “Cluster 4”, “Cluster 1”. For batch 2 (Supplemental Fig. S6), the ArchR object was clustered at a resolution of 0.2 and the “addTrajectory” function used with the following supervised trajectory: “Cluster 3”, “Cluster 4”, “Cluster 5”. Transcription factor gene expression and corresponding motif accessibility (ChromVAR deviation scores) were correlated over pseudotime using “getTrajectory” and “correlateTrajectories” using “varCutOff = 0.55” for both RNA and motif matrices. Transcription factors were filtered according to correlation false discovery rate < 0.001, and fold enrichment of motif accessibility in epithelial over stromal lineage as determined by the “peakAnnoEnrichment” function >1. Transcription factors were further filtered according to correlation value, with a value of >0.2 used to identify activators, and <-0.2 to identify repressors. Activators or repressors were ordered according to gene expression along pseudotime, and the R package pheatmap (v1.0.12) was used to generate paired heatmaps of gene expression and corresponding motif accessibility along pseudotime (Figure 5). Gene expression and motif accessibility (as ChromVar deviations) were visualised along pseudotime using the “plotTrajectory” function.

## QUANTIFICATION AND STATISTICAL ANALYSIS

All the details of quantifications and statistical analyses are fully described in the main text, figure legends, and METHOD DETAILS section.

### Supplemental Tables

**Supplemental Table 1.** Marker genes for clusters in day 14 epithelium corresponding to nephron segments (from batch 3, as in Supplemental Fig. S5E), generated using snRNA-seq data (Wilcoxon test, FDR ≤ 0.01 & Log2FC ≥ 1).

**Supplemental Table 2.** Marker genes for clusters representing epithelial lineage time point, and stromal lineage all time points combined (from batch 3, as in Main Fig. 1E), generated using snRNA-seq data (FDR ≤ 0.01 & Log2FC ≥ 1).

**Supplemental Table 3.** Gene Ontology (GO) analysis of marker genes from Supplemental Table 2 (epithelial lineage only)

**Supplemental Table 4.** Marker genes for clusters representing epithelial or stromal lineage (from batch 3, as in Main Fig. 2A), generated using snRNA-seq data (FDR ≤ 0.01 & Log2FC ≥ 1).

**Supplemental Table 5.** Gene Ontology (GO) analysis of the PAX8 regulon

**Supplemental Table 6.** Marker genes for clusters corresponding to DMSO-treated or CHIR-treated epithelium at day 14 (from batch 3, as in Main Fig. 7F & I), generated using snRNA-seq data (FDR ≤ 0.01 & Log2FC ≥ 1).

**Supplemental Table 7.** Gene Ontology (GO) analysis of the genes enriched in DMSO-treated epithelium compared to CHIR-treated epithelium at day 14 (from Supplemental Table 6)

**Supplemental Table 8.** Peak to gene links, generated with ArchR (correlation > 0.4, FDR < 1e-4). Values in ‘Strand’ column are arbitrary and included to make the table compatible with the ‘findMotifsGenome.pl’ function in HOMER.

## Supplemental Figure Legends

**Supplemental Figure S1, related to Figure 1.**
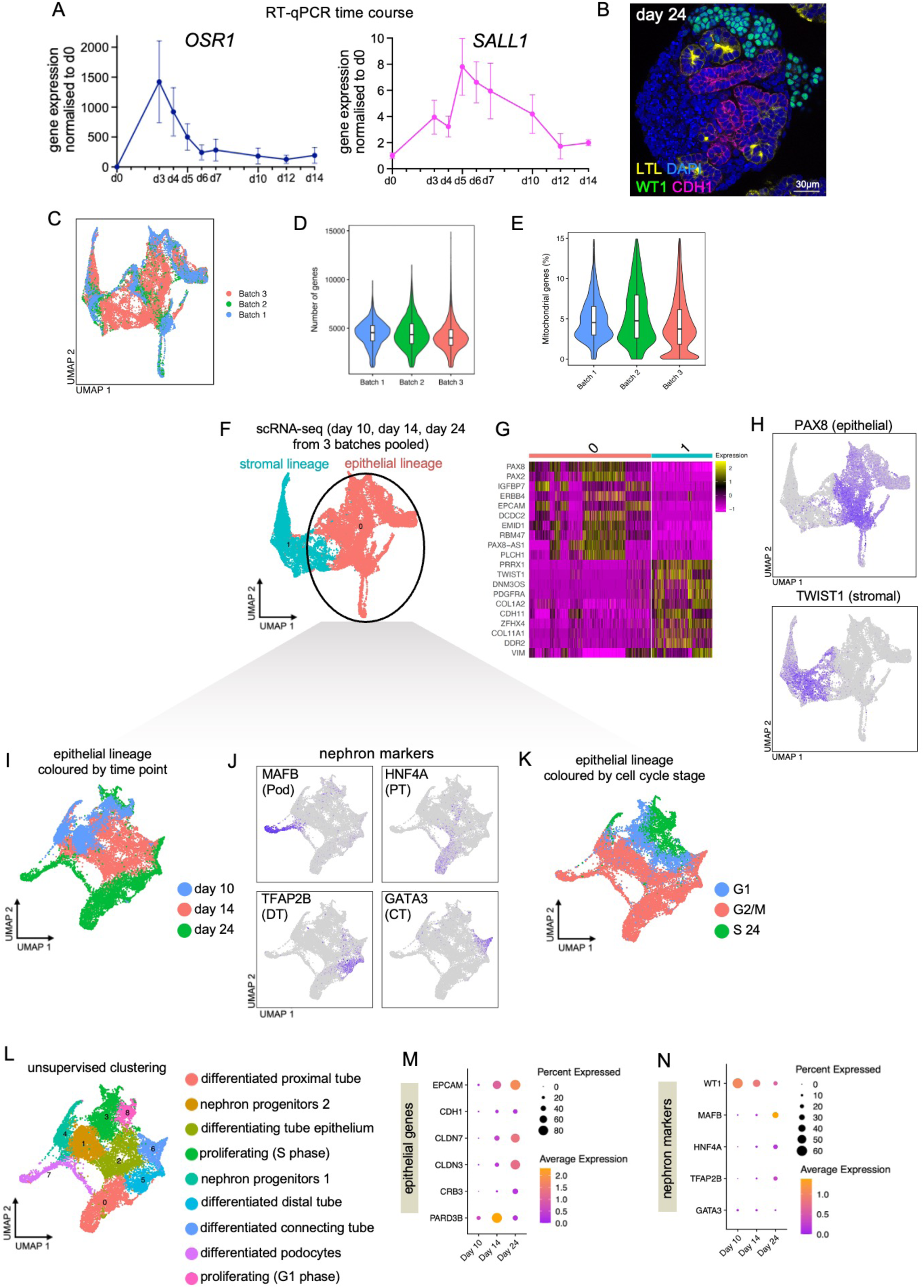
Single cell RNA-seq characterisation of renal organoids. (A) Time course RT-qPCR data for *OSR1* and *SALL1* over the period of the kidney organoid protocol. Dots and bars represent mean ± SEM, N = 2-4 independent organoid batches at each time point. (B) Representative immunofluorescence image of a renal organoid at day 24 stained for WT1 (green), LTL (white), CDH1 (red) and DAPI (blue). Scale bar is 30µm. (C) UMAP plot of 23,856 sc-RNA-seq and sn-RNA-seq cells harvested across 3 organoid batches, harvested at days 10, 14 and 24, coloured by batch. (D) Violin plots showing number of genes per batch. (E) Violin plots showing percentage mitochondrial reads per batch. Box center line, median; limits, upper and lower quartiles; whiskers, 1.5x interquartile range (F) UMAP plot of 23,856 sc-RNA-seq and sn-RNA-seq cells as in (C), coloured by epithelial or stromal lineage. (G) Heatmap showing top 10 genes that define the epithelial and stromal lineages. (H) UMAP plots of 23,856 sc-RNA-seq and sn-RNA-seq cells as in (C), coloured by *PAX8* expression (top) and *TWIST1* expression (bottom). (I) UMAP plot of 15,917 sc-RNA-seq and sn-RNA-seq cells corresponding to the epithelial lineage as in (F), re-projected and coloured according to time point. (J) UMAP plots of 15,917 6 sc-RNA-seq and sn-RNA-seq cells as in (I), coloured by *MAFB* expression (top left), *HNF4A* expression (top right), *TFAP2B* expression (bottom left) and *GATA3* expression (bottom right). (K) UMAP plots of 15,917 6 sc-RNA-seq and sn-RNA-seq cells as in (I), coloured by cell cycle phase. (L) UMAP plots of 15,917 6 sc-RNA-seq and sn-RNA-seq cells as in (I), coloured by unsupervised clustering, annotated according to querying marker genes for each cluster with the Human Nephrogenesis Atlas (https://sckidney.flatironinstitute.org) (Lindstrom et al., 2021) and assessment of cell cycle phase. (M) Dot plot of gene expression for epithelial genes in 15,917 6 sc-RNA-seq and sn-RNA-seq cells as in (D) separated by time point. (N) Dot plot of gene expression for nephron marker genes in 15,917 6 sc-RNA-seq and sn-RNA-seq cells as in (D) separated by time point.

**Supplemental Figure S2, related to Figures 1 and 3.**
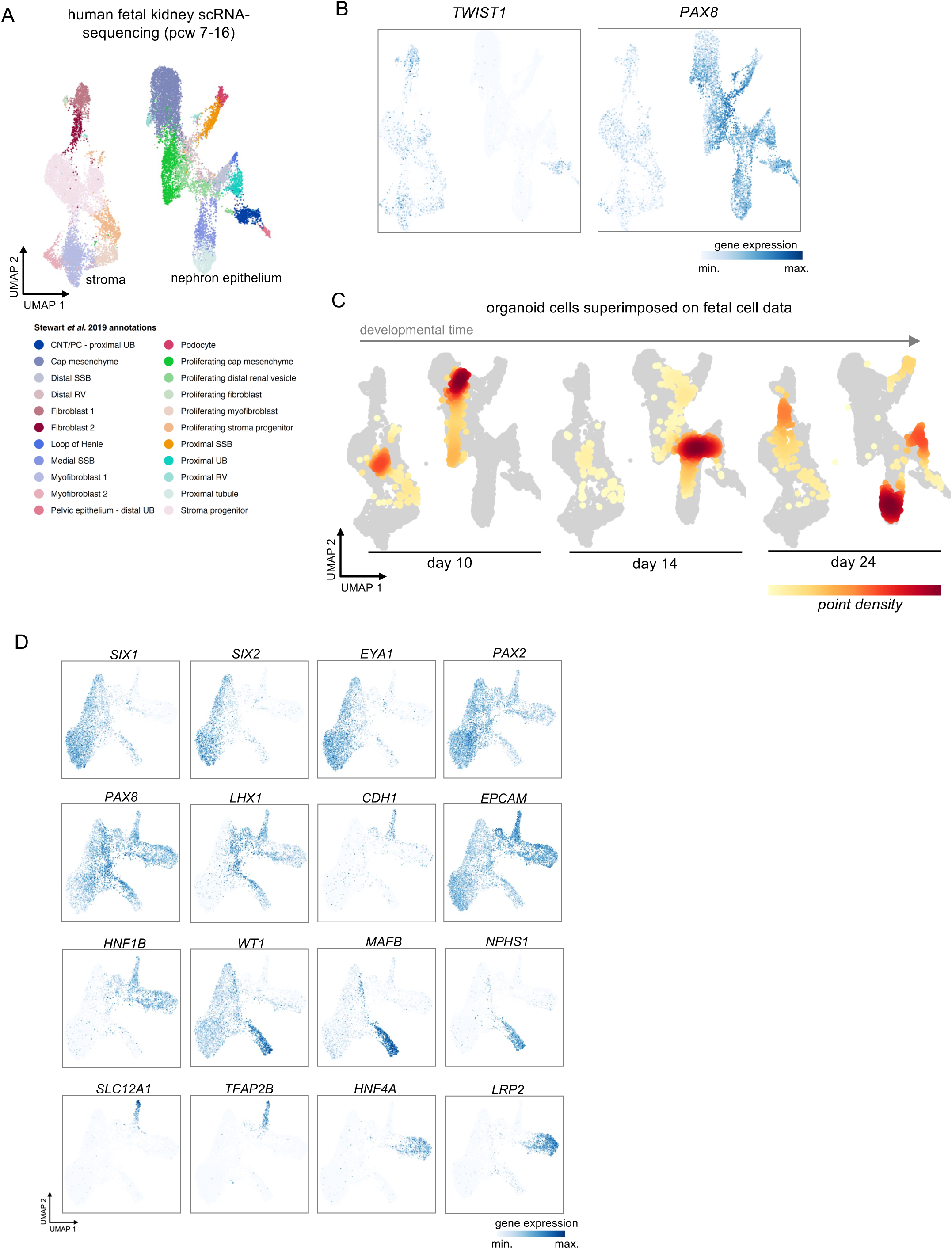
Comparison of single cell RNA-seq data of human renal organoids to human fetal kidney *in vivo*. (A) UMAP plot of 17,994 sc-RNA-seq cells from 6 human fetal kidney samples collected between post conception week (pcw) 7 and 16, split by epithelial and stromal lineages, coloured by cell type annotations (Stewart et al., 2019). (B) UMAP plot of 17,994 sc-RNA-seq cells as in (A), coloured by gene expression of the stromal marker *TWIST1* (left) or the nephron marker *PAX8* (right). (C) Asymmetric integration of kidney organoid sc-RNA-seq data split by time point with human fetal kidney sc-RNA-seq data, coloured by point density of organoid data. (D) UMAP plots of 8,862 nephron epithelial cells from (A), isolated and re-projected, and coloured by gene expression of nephron marker genes (related to Figure 3).

**Supplemental Figure S3, related to Figures 1 and 2.**
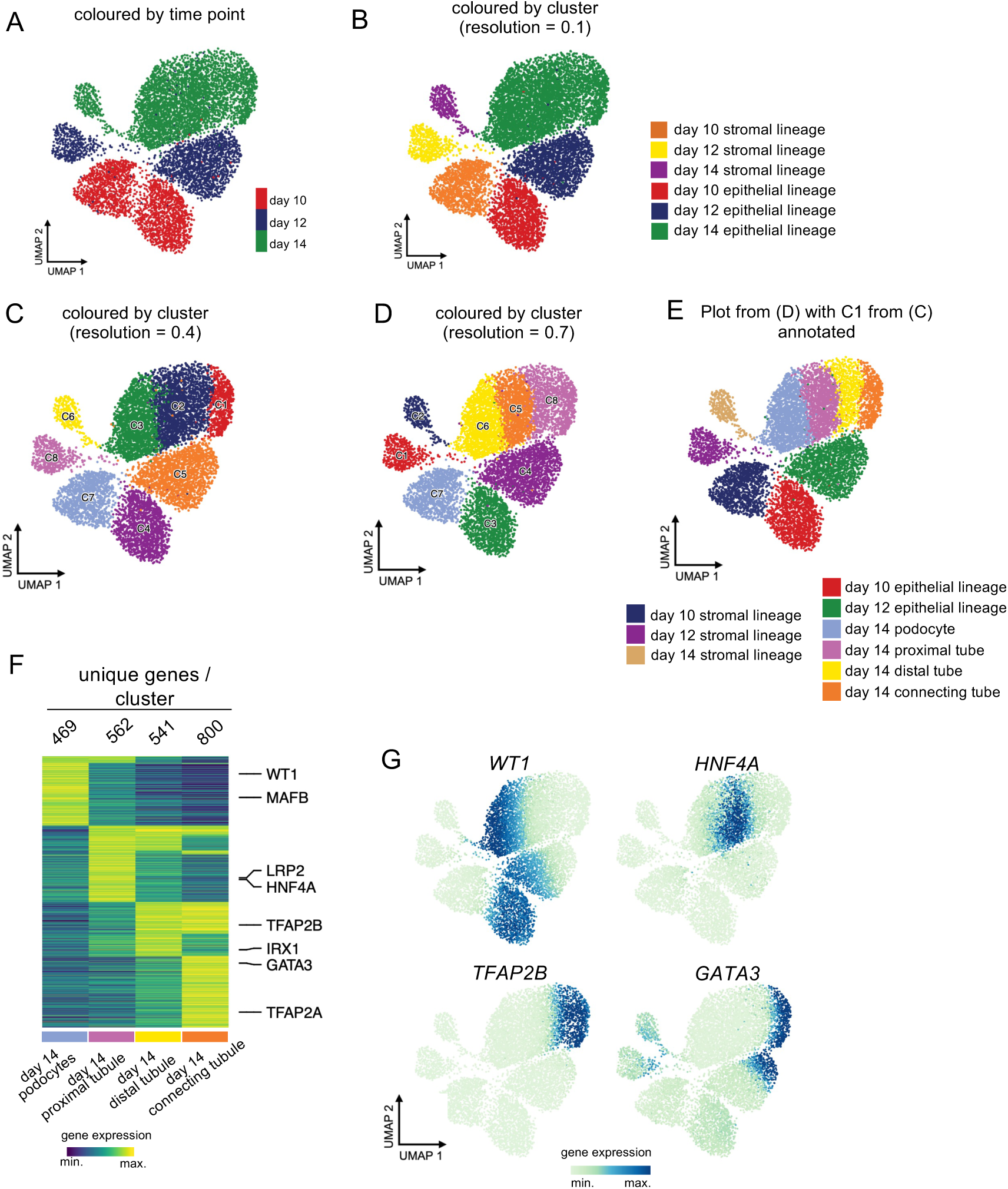
Clustering multi-ome cells reveals nephron cell types in day 14 epithelium. (A) UMAP plot of 9,147 multi-ome cells from batch 3 coloured by time point. (B) UMAP plot of 9,147 multi-ome cells from batch 3 coloured by unsupervised clustering (resolution 0.1) and annotated according to time point and epithelial or stromal lineage. (C) UMAP plot of 9,147 multi-ome cells from batch 3 coloured by unsupervised clustering (resolution 0.4). (D) UMAP plot of 9,147 multi-ome cells from batch 3 coloured by unsupervised clustering (resolution 0.7). (E) UMAP plot of 9,147 multi-ome cells from batch 3 coloured by unsupervised clustering (resolution 0.7), annotated according to time point, lineage and cell type as determined by marker genes identified from differential gene expression analysis (in E, F), with cells corresponding to C1 in (C) labelled as connecting tube. (F) Heat map of expression of marker genes for clusters in (D) determined by snRNA-seq with representative markers annotated (left, log2FC > 1, FDR < 0.01, two-sided Wilcoxon rank-sum test; log2FC: log2 fold change; FDR: false discovery rate). (G) UMAP plots of 9,147 multi-ome cells from batch 3 coloured by *WT1*, *HNF4A*, *TFAP2B* or *GATA3* gene expression.

**Supplemental Figure S4, related to Figure 2.**
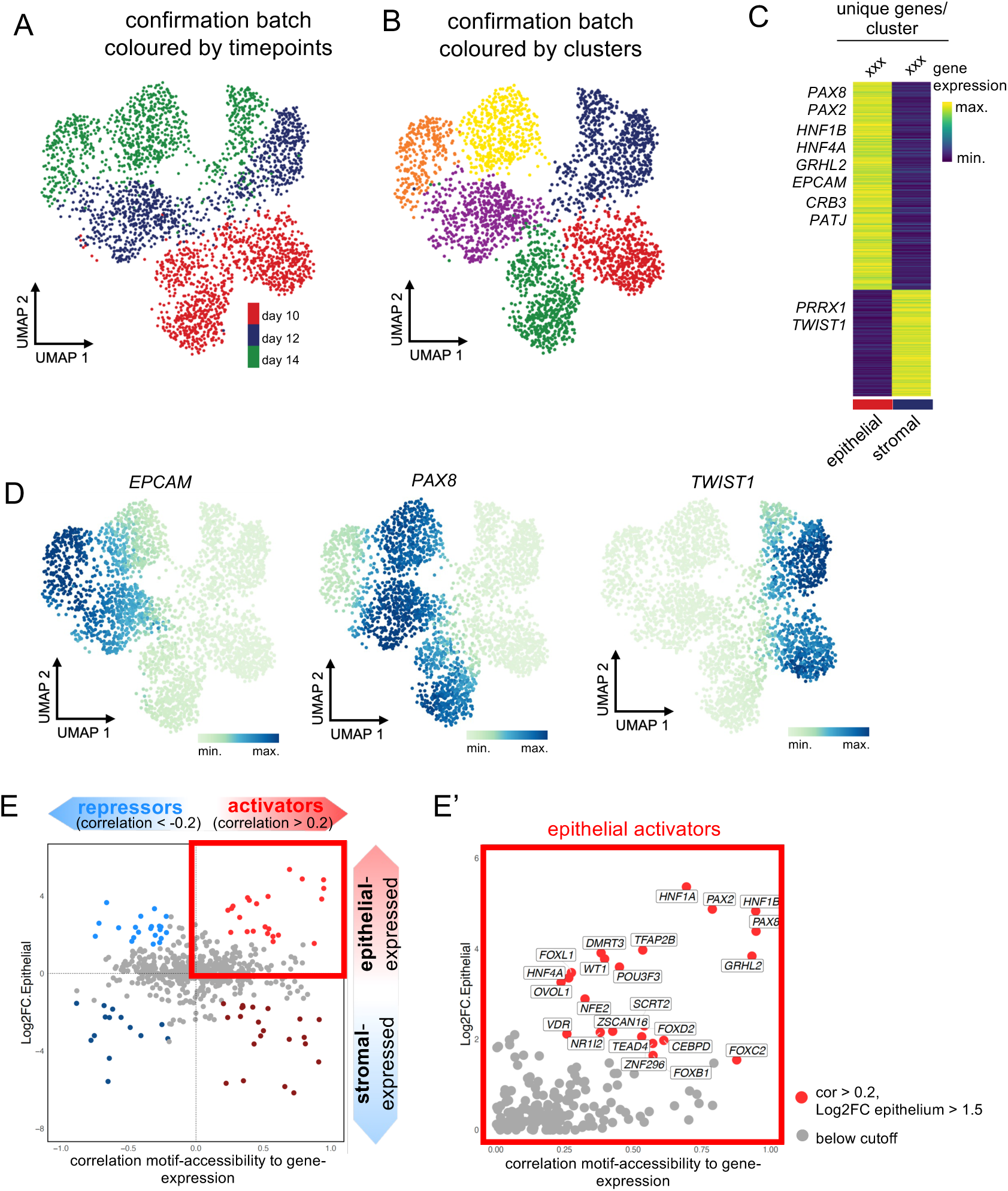
Analysis of an independent organoid multi-ome batch. (A) UMAP of 3,040 cells from batch 2 clustered according to time point. (B) UMAP of 3,040 cells from batch 2 clustered according to unsupervised clustering. (C) Heat map of expression of marker genes for clusters in (A) determined by snRNA-seq with representative markers annotated (log2 FC > 1, FDR < 0.01, two-sided Wilcoxon rank-sum test). (D) Heat map of marker peaks of accessible chromatin for clusters in (A) determined by snATAC-seq (left, log2 FC > 1, FDR < 0.01, two-sided Wilcoxon rank-sum test). (E, E’) Transcription factors plotted according to Pearson correlation coefficient (PCC) of gene expression versus corresponding motif accessibility, and log2FC gene expression in the epithelial compared to the stromal lineage as determined by snRNA-seq. Thresholds used to colour points according to principle outlined in (Main Figure 2D’): PCC > 0.2, Log2FC epithelial lineage > 1.5 (red = activators), PCC < -0.2, Log2FC epithelial lineage > 1.5 (blue = repressors).

**Supplemental Figure S5, related to Figures 4 and 5.**
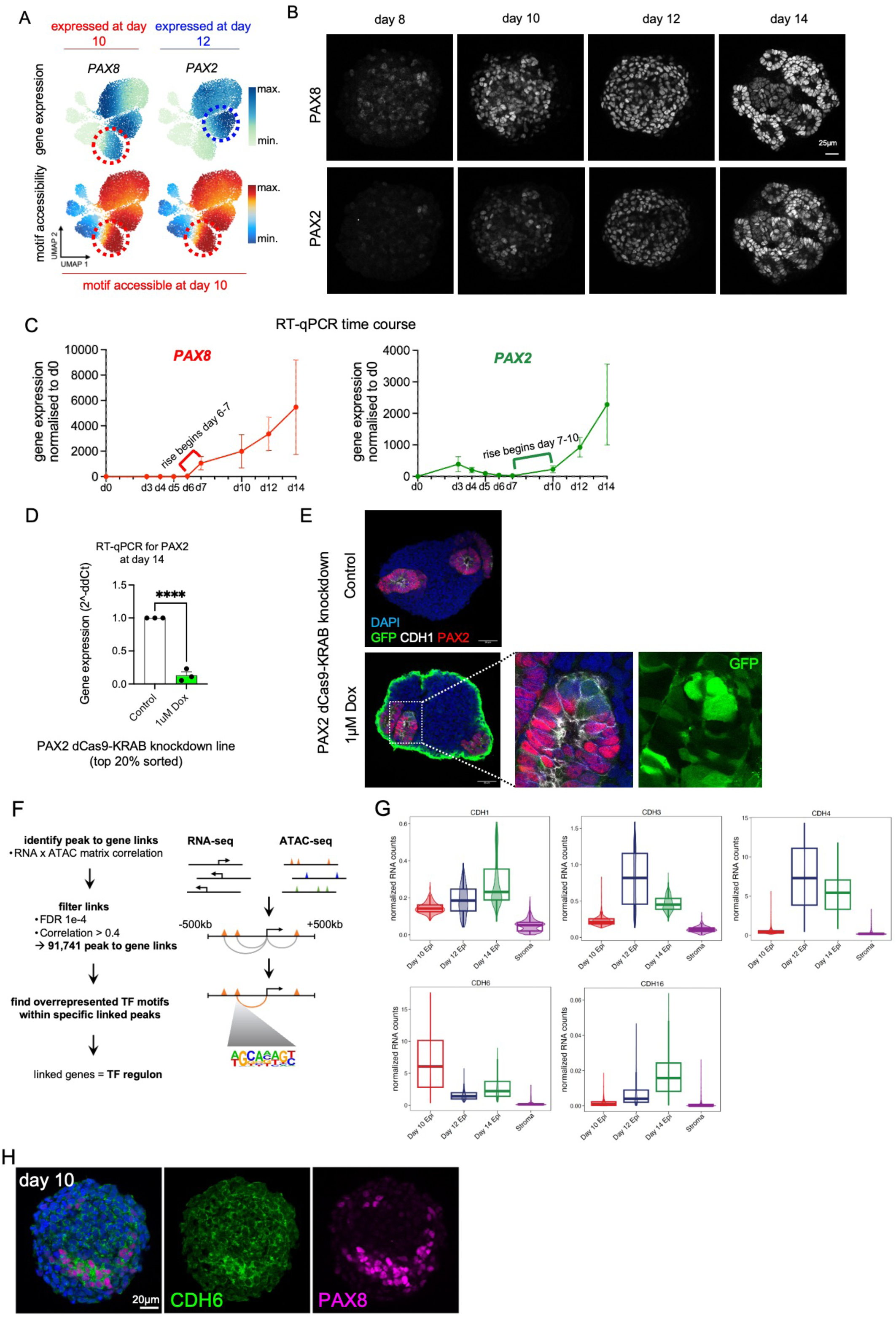
Expression of PAX2 and PAX8 in human renal organoid MET, and identifying the PAX8 regulon. (A) UMAP plots of 9,147 multi-ome cells from batch 3 showing gene expression (top) and motif accessibility (bottom) for PAX8 and PAX2, highlighting start of expression of PAX8 at day 10 and of PAX2 at day 12, but accessibility of the shared motif apparent at day 10. (B) Immunofluorescence analysis of representative organoids at days 8, 10, 12 and 14, stained for PAX8 (top) or PAX2 (bottom), scale bars 25µm. (C) Time course RT-qPCR data for *PAX8* and *PAX2* over the period of the kidney organoid protocol. Dots and bars represent mean ± SEM, N = 2-4 independent organoid batches at each time point. (D) RT-qPCR for *PAX2* in organoids generated from PAX2-dCas9-KRAB iPSCs (top 20% sorted, see STAR METHODS) and harvested at day 14, either with no treatment (control, white bars) or following Dox treatment from day 4 (1μM Dox, green bars). N = 3 independent organoid batches. Dots represent data from each batch normalised to corresponding control, with bars representing mean ± SEM. Unpaired t-test, **** (p < 0.0001), compared to control. (E) Immunofluorescence of organoids generated from PAX2-dCas9-KRAB iPSCs and harvested at day 14, either with no treatment (control, top) or following Dox treatment from day 4 (1μM Dox, bottom), showing GFP (green), PAX2 (magenta), E-Cadherin/CDH1 (white) with DAPI as counterstain (blue, nuclei). Scale bars 25µm. (F) Schematic of strategy used to identify peak to gene links (related to Figure 4). (G) Violin plots of gene expression of *CDH1*, *CDH3, CDH4, CDH6* and *CDH16* in 9,147 multi-ome cells from batch 3 split by time point (related to Figure 4). (H) Immunofluorescence analysis of representative organoids at day 10 showing co-localisation of PAX8 (white) and CDH6 (green). Scale bars 20µm (related to Figure 4).

**Supplemental Figure S6, related to Figure 6.**
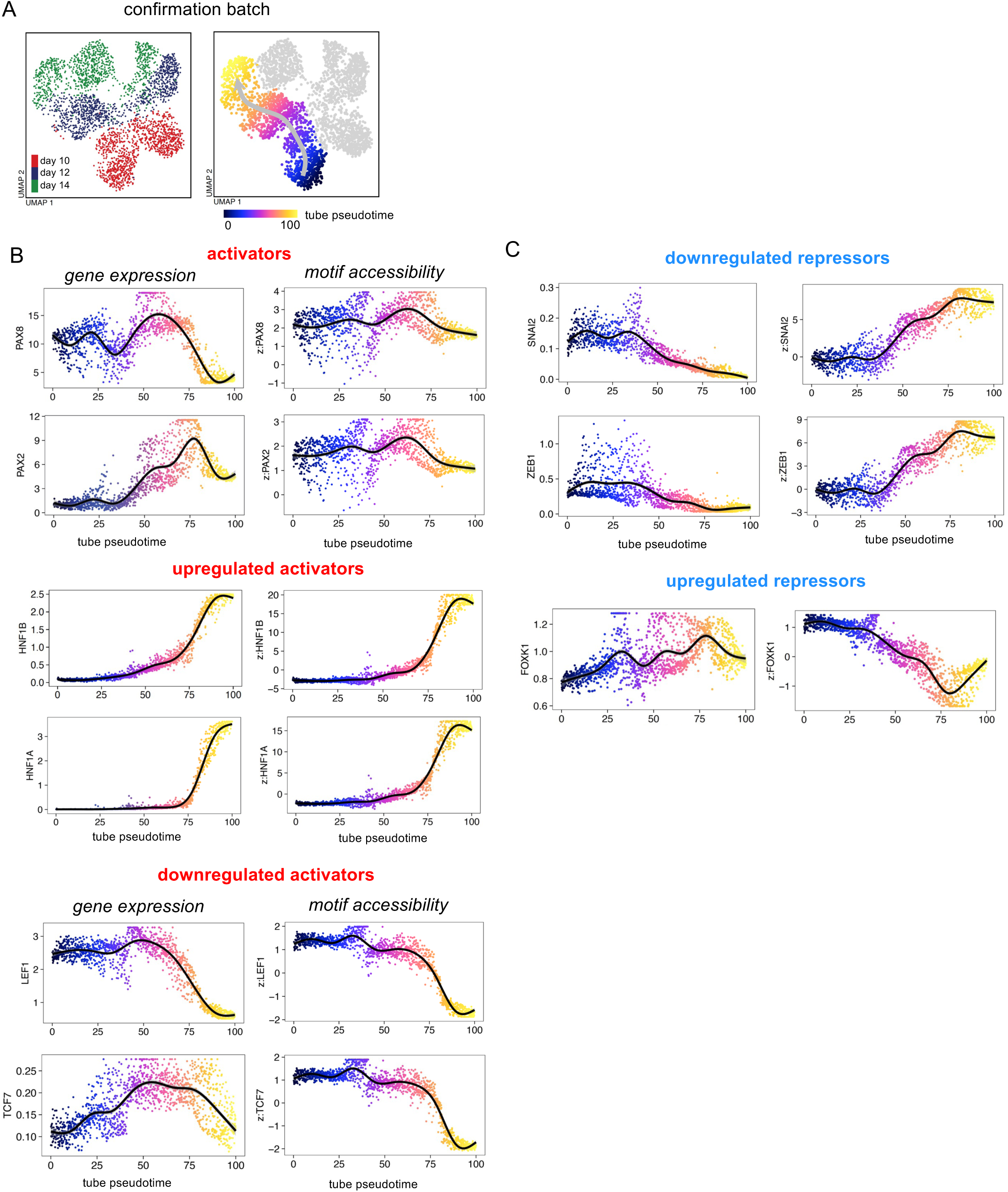
Transcription factor dynamics in an independent organoid multi-ome batch. (A) UMAP plot of 3,040 cells from batch 2 coloured according to tube epithelial pseudotime trajectory, generated in ArchR (Granja et al., 2021). (B) Cells in batch 2 plotted according to tube epithelial pseudotime as in (A), and gene expression (left) or motif accessibility (right), of the transcriptional activators *PAX8, PAX2* which show consistent motif accessibility along pseudotime, an example upregulated activator *HNF1B,* and example downregulated activators *LEF1* and *TCF7*. (C) Cells in batch 2 plotted according to tube epithelial pseudotime as in (A), and gene expression (left) or motif accessibility (right), of example downregulated transcriptional repressors *SNAI2,* and *ZEB1,* and example upregulated repressor *FOXK1*.

**Supplemental Figure S7, related to Figure 7.**
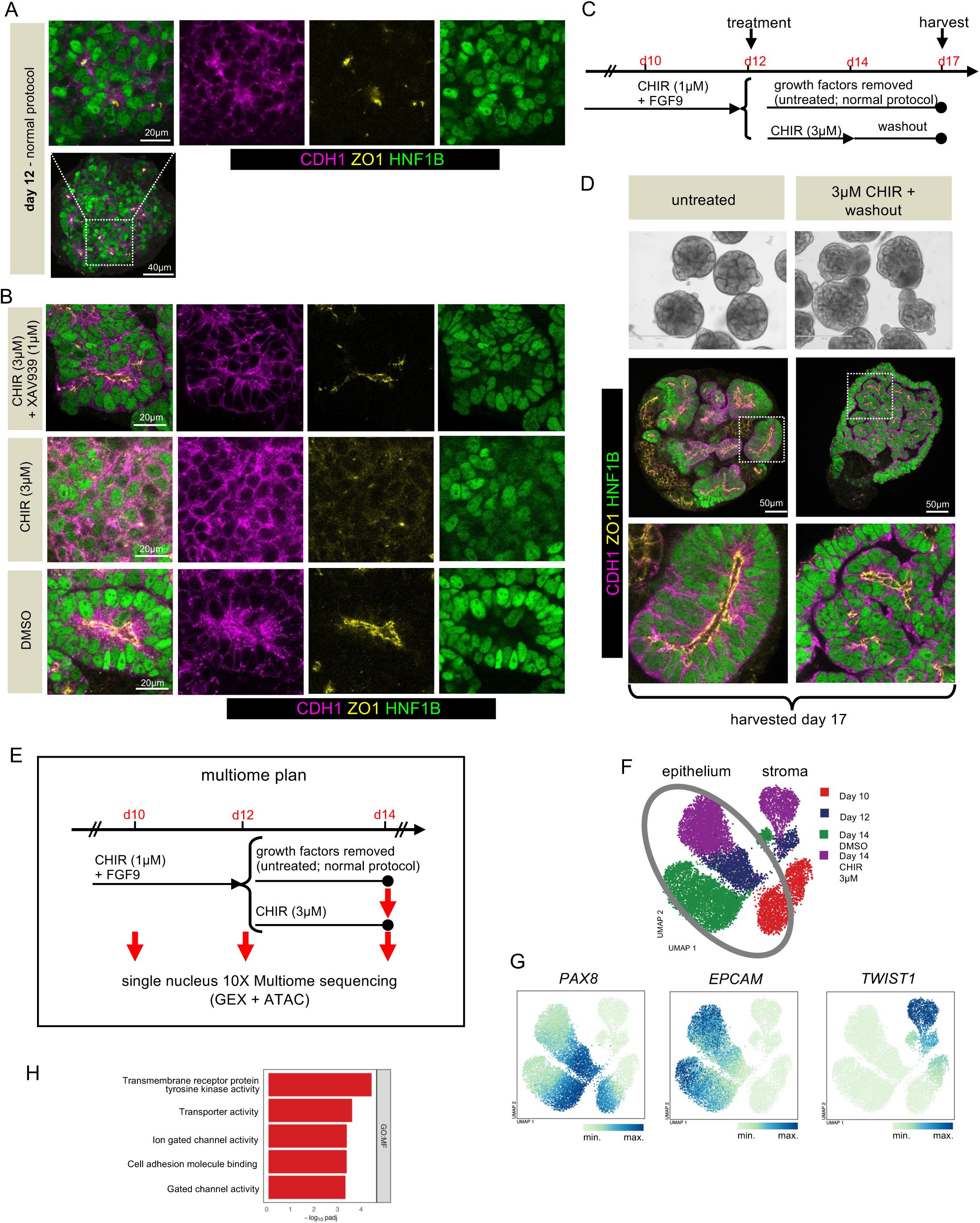
Effect of persistent Wnt/β-Catenin activation on MET in human renal organoids. (A) Immunofluorescence of organoids at day 12 from same batch as shown in (B) and in Main Fig. 7, showing expression of CDH1 (magenta), ZO1 (yellow) and HNF1B (green). Scale bars 20µm (top), 40µm (bottom). (B) Insets from Fig. 7 of organoids treated with the indicated conditions between days 12 to 14, to show individual channels of CDH1 (magenta), ZO1 (yellow) and HNF1B (green). Scale bars 20µm. (C) Schematic of experiment to test reversibility of effect of sustained Wnt/β-Catenin signalling with CHIR washout between days 14 and 17. (D) Organoids harvested at day 17 following treatments indicated in (A), with light microscopy images (top, scale bar 400µm), and immunofluorescence images showing expression of HNF1B (green), E-Cadherin/CDH1 (magenta) and ZO1 (yellow). Dotted white boxes indicate positions of magnification panels below. Scale bars 50µm. (E) Schematic of renal organoid multi-ome profiling strategy for batch 3 – corresponding to schematic in Main Fig. 1D but including including the two day 14 samples sequenced in parallel: one treated with DMSO from day 12 to 14 (analogous to normal protocol), and one treated with 3µM CHIR from day 12 to 14. (F) UMAP plot of 13,502 multi-ome cells from batch 3, of organoids harvested at day 10, day 12, and at day 14 treated either with DMSO or 3µM CHIR from day 12 to 14 (See Fig. 1D), coloured according to sample. (G) UMAP plots of 13,502 multi-ome cells from batch 3 as in (D), coloured according to gene expression of *PAX8* (left)*, EPCAM* (centre), or *TWIST1* (right). (H) Gene ontology analysis of genes enriched in day 14 epithelium treated with DMSO from day 12-14 compared to 3µM CHIR, showing top 5 highest specific GO Molecular Function terms by log10 adjusted p value, full list in Supplemental Table 7.

**Supplemental Figure S8, related to Figure 7.**
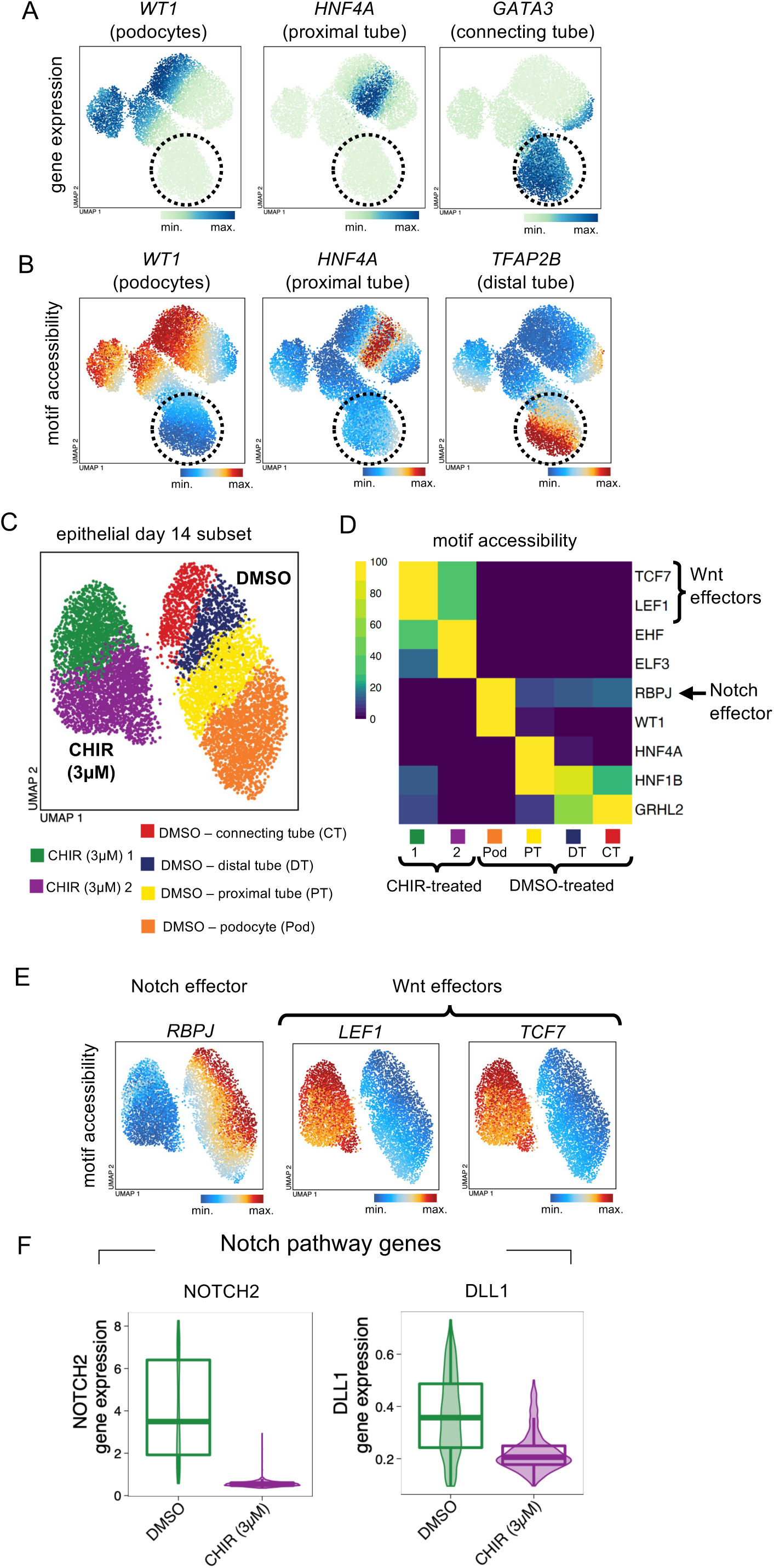
Persistent Wnt/μ-Catenin activation in human renal organoids distalises the epithelium via inhibition of Notch signalling. (A) UMAP plots of 10,793 multi-ome cells from batch 3 (as in Main Fig. 7C) showing altered gene expression of *WT1, HNF4A,* and *GATA3* in day 14 cells treated with 3µM CHIR from day 12 to 14 (highlighted by dotted circle) compared to day 14 cells cultured according to the normal protocol. (B) UMAP plots of multi-ome cells from batch 3 (as in Main Fig. 7C) showing altered motif accessibility of WT1, HNF4A, and TFAP2B in day 14 cells treated with 3µM CHIR from day 12 to 14 (highlighted by dotted circle) compared to day 14 cells cultured according to the normal protocol. (C) UMAP plot of 7,871 multi-ome cells from batch 3 comprising only day 14 cells treated either with DMSO or 3µM CHIR from day 12 to 14, subjected to unsupervised clustering and coloured according to nephron segment (in DMSO condition). The CHIR-treated epithelium showed 2 clusters by unsupervised clustering. (D) Heatmap of motif accessibility of selected transcription factors in cell clusters as in (C). Accessibility of TCF7, LEF1, EHF and ELF3 motifs is increased in day 14 epithelium treated with 3µM CHIR from day 12 to 14, consistent with sustained Wnt/b-Catenin activity. Accessibility of RBPJ motifs, and of WT1 and HNF4A motifs, are lost in CHIR-treated epithelium, consistent with inhibition of Notch signalling and a loss of proximal identity, respectively. HNF1B and GRHL2 motifs are greatly reduced in CHIR-treated epithelium. (E) UMAP plots of multi-ome cells from batch 3 (as in C) showing altered motif accessibility of RBPJ, LEF1 and TCF7 in day 14 cells treated with 3µM CHIR from day 12 to 14 compared to day 14 cells cultured according to the normal protocol. (F) Violin plots of multi-ome cells from batch 3 (as in C) showing reduced gene expression of the Notch pathway receptor *NOTCH2* and of the Notch ligand *DLL1* expression in CHIR-treated cells at day 14 compared to day 14 cells cultured according to the normal protocol. Box center line, median; limits, upper and lower quartiles; whiskers, 1.5x interquartile range.

## References

1. Adam, M., Potter, A. S., & Potter, S. S. (2017). Psychrophilic proteases dramatically reduce single-cell RNA-seq artifacts: a molecular atlas of kidney development. Development, 144(19), 3625–3632. https://doi.org/10.1242/dev.151142

2. Alotaibi, H., Basilicata, M. F., Shehwana, H., Kosowan, T., Schreck, I., Braeutigam, C., Konu, O., Brabletz, T., & Stemmler, M. P. (2015). Enhancer cooperativity as a novel mechanism underlying the transcriptional regulation of E-cadherin during mesenchymal to epithelial transition. Biochim Biophys Acta, 1849(6), 731–742. https://doi.org/10.1016/j.bbagrm.2015.01.005

3. Alpert, A., Moore, L. S., Dubovik, T., & Shen-Orr, S. S. (2018). Alignment of single-cell trajectories to compare cellular expression dynamics. Nat Methods, 15(4), 267–270. https://doi.org/10.1038/nmeth.4628

4. Azzolin, L., Panciera, T., Soligo, S., Enzo, E., Bicciato, S., Dupont, S., Bresolin, S., Frasson, C., Basso, G., Guzzardo, V., Fassina, A., Cordenonsi, M., & Piccolo, S. (2014). YAP/TAZ incorporation in the beta-catenin destruction complex orchestrates the Wnt response. Cell, 158(1), 157–170. https://doi.org/10.1016/j.cell.2014.06.013

5. Bleu, M., Mermet-Meillon, F., Apfel, V., Barys, L., Holzer, L., Bachmann Salvy, M., Lopes, R., Amorim Monteiro Barbosa, I., Delmas, C., Hinniger, A., Chau, S., Kaufmann, M., Haenni, S., Berneiser, K., Wahle, M., Moravec, I., Vissieres, A., Poetsch, T., Ahrne, E., . . . Galli, G. G. (2021). PAX8 and MECOM are interaction partners driving ovarian cancer. Nat Commun, 12(1), 2442. https://doi.org/10.1038/s41467-021-22708-w

6. Bolos, V., Peinado, H., Perez-Moreno, M. A., Fraga, M. F., Esteller, M., & Cano, A. (2003). The transcription factor Slug represses E-cadherin expression and induces epithelial to mesenchymal transitions: a comparison with Snail and E47 repressors. J Cell Sci, 116(Pt 3), 499–511. https://doi.org/10.1242/jcs.00224

7. Bouchard, M., Pfeffer, P., & Busslinger, M. (2000). Functional equivalence of the transcription factors Pax2 and Pax5 in mouse development. Development, 127(17), 3703–3713. https://doi.org/10.1242/dev.127.17.3703

8. Bouchard, M., Souabni, A., Mandler, M., Neubuser, A., & Busslinger, M. (2002). Nephric lineage specification by Pax2 and Pax8. Genes Dev, 16(22), 2958–2970. https://doi.org/10.1101/gad.240102

9. Boutet, A., De Frutos, C. A., Maxwell, P. H., Mayol, M. J., Romero, J., & Nieto, M. A. (2006). Snail activation disrupts tissue homeostasis and induces fibrosis in the adult kidney. EMBO J, 25(23), 5603–5613. https://doi.org/10.1038/sj.emboj.7601421

10. Bowman, C. J., Ayer, D. E., & Dynlacht, B. D. (2014). Foxk proteins repress the initiation of starvation-induced atrophy and autophagy programs. Nat Cell Biol, 16(12), 1202–1214. https://doi.org/10.1038/ncb3062

11. Brakeman, P. R., Liu, K. D., Shimizu, K., Takai, Y., & Mostov, K. E. (2009). Nectin proteins are expressed at early stages of nephrogenesis and play a role in renal epithelial cell morphogenesis. Am J Physiol Renal Physiol, 296(3), F564–574. https://doi.org/10.1152/ajprenal.90328.2008

12. Campbell, K. R., & Yau, C. (2017). switchde: inference of switch-like differential expression along single-cell trajectories. Bioinformatics, 33(8), 1241–1242. https://doi.org/10.1093/bioinformatics/btw798

13. Carroll, T. J., Park, J. S., Hayashi, S., Majumdar, A., & McMahon, A. P. (2005). Wnt9b plays a central role in the regulation of mesenchymal to epithelial transitions underlying organogenesis of the mammalian urogenital system. Dev Cell, 9(2), 283–292. https://doi.org/10.1016/j.devcel.2005.05.016

14. Carvalho, A., Hermanns, P., Rodrigues, A. L., Sousa, I., Anselmo, J., Bikker, H., Cabral, R., Pereira-Duarte, C., Mota-Vieira, L., & Pohlenz, J. (2013). A new PAX8 mutation causing congenital hypothyroidism in three generations of a family is associated with abnormalities in the urogenital tract. Thyroid, 23(9), 1074–1078. https://doi.org/10.1089/thy.2012.0649

15. Chan, S. C., Hajarnis, S. S., Vrba, S. M., Patel, V., & Igarashi, P. (2020). Hepatocyte nuclear factor 1beta suppresses canonical Wnt signaling through transcriptional repression of lymphoid enhancer-binding factor 1. J Biol Chem, 295(51), 17560–17572. https://doi.org/10.1074/jbc.RA120.015592

16. Chan, S. C., Zhang, Y., Pontoglio, M., & Igarashi, P. (2019). Hepatocyte nuclear factor-1beta regulates Wnt signaling through genome-wide competition with beta-catenin/lymphoid enhancer binding factor. Proc Natl Acad Sci U S A, 116(48), 24133–24142. https://doi.org/10.1073/pnas.1909452116

17. Charlton, J. R., Baldelomar, E. J., Hyatt, D. M., & Bennett, K. M. (2021). Nephron number and its determinants: a 2020 update. Pediatr Nephrol, 36(4), 797–807. https://doi.org/10.1007/s00467-020-04534-2

18. Corces, M. R., Granja, J. M., Shams, S., Louie, B. H., Seoane, J. A., Zhou, W., Silva, T. C., Groeneveld, C., Wong, C. K., Cho, S. W., Satpathy, A. T., Mumbach, M. R., Hoadley, K. A., Robertson, A. G., Sheffield, N. C., Felau, I., Castro, M. A. A., Berman, B. P., Staudt, L. M., . . . Chang, H.Y. (2018). The chromatin accessibility landscape of primary human cancers. Science, 362(6413). https://doi.org/10.1126/science.aav1898

19. Costantini, F., & Kopan, R. (2010). Patterning a complex organ: branching morphogenesis and nephron segmentation in kidney development. Dev Cell, 18(5), 698–712. https://doi.org/10.1016/j.devcel.2010.04.008

20. Dutta, S., Mana-Capelli, S., Paramasivam, M., Dasgupta, I., Cirka, H., Billiar, K., & McCollum, D. (2018). TRIP6 inhibits Hippo signaling in response to tension at adherens junctions. EMBO Rep, 19(2), 337–350. https://doi.org/10.15252/embr.201744777

21. Duvall, K., Crist, L., Perl, A. J., Pode Shakked, N., Chaturvedi, P., & Kopan, R. (2022). Revisiting the role of Notch in nephron segmentation confirms a role for proximal fate selection during mouse and human nephrogenesis. Development, 149(10). https://doi.org/10.1242/dev.200446

22. Eger, A., Aigner, K., Sonderegger, S., Dampier, B., Oehler, S., Schreiber, M., Berx, G., Cano, A., Beug, H., & Foisner, R. (2005). DeltaEF1 is a transcriptional repressor of E-cadherin and regulates epithelial plasticity in breast cancer cells. Oncogene, 24(14), 2375–2385. https://doi.org/10.1038/sj.onc.1208429

23. Elias, K. M., Emori, M. M., Westerling, T., Long, H., Budina-Kolomets, A., Li, F., MacDuffie, E., Davis, M. R., Holman, A., Lawney, B., Freedman, M. L., Quackenbush, J., Brown, M., & Drapkin, R. (2016). Epigenetic remodeling regulates transcriptional changes between ovarian cancer and benign precursors. JCI Insight, 1(13). https://doi.org/10.1172/jci.insight.87988

24. Faure, L., Soldatov, R., Kharchenko, P. V., & Adameyko, I. (2023). scFates: a scalable python package for advanced pseudotime and bifurcation analysis from single-cell data. Bioinformatics, 39(1). https://doi.org/10.1093/bioinformatics/btac746

25. Ferre, S., & Igarashi, P. (2019). New insights into the role of HNF-1beta in kidney (patho)physiology. Pediatr Nephrol, 34(8), 1325–1335. https://doi.org/10.1007/s00467-018-3990-7

26. Fornes, O., Castro-Mondragon, J. A., Khan, A., van der Lee, R., Zhang, X., Richmond, P. A., Modi, B. P., Correard, S., Gheorghe, M., Baranasic, D., Santana-Garcia, W., Tan, G., Cheneby, J., Ballester, B., Parcy, F., Sandelin, A., Lenhard, B., Wasserman, W. W., & Mathelier, A. (2020). JASPAR 2020: update of the open-access database of transcription factor binding profiles. Nucleic Acids Res, 48(D1), D87–D92. https://doi.org/10.1093/nar/gkz1001

27. Gilbert, L. A., Horlbeck, M. A., Adamson, B., Villalta, J. E., Chen, Y., Whitehead, E. H., Guimaraes, C., Panning, B., Ploegh, H. L., Bassik, M. C., Qi, L. S., Kampmann, M., & Weissman, J. S. (2014). Genome-Scale CRISPR-Mediated Control of Gene Repression and Activation. Cell, 159(3), 647–661. https://doi.org/10.1016/j.cell.2014.09.029

28. Gilmour, D., Rembold, M., & Leptin, M. (2017). From morphogen to morphogenesis and back. Nature, 541(7637), 311–320. https://doi.org/10.1038/nature21348

29. Gong, K. Q., Yallowitz, A. R., Sun, H., Dressler, G. R., & Wellik, D. M. (2007). A Hox-Eya-Pax complex regulates early kidney developmental gene expression. Mol Cell Biol, 27(21), 7661–7668. https://doi.org/10.1128/MCB.00465-07

30. Gonzalez, D. M., & Medici, D. (2014). Signaling mechanisms of the epithelial-mesenchymal transition. Sci Signal, 7(344), re8. https://doi.org/10.1126/scisignal.2005189

31. Granja, J. M., Corces, M. R., Pierce, S. E., Bagdatli, S. T., Choudhry, H., Chang, H. Y., & Greenleaf, W. J. (2021). ArchR is a scalable software package for integrative single-cell chromatin accessibility analysis. Nat Genet, 53(3), 403–411. https://doi.org/10.1038/s41588-021-00790-6

32. Heinz, S., Benner, C., Spann, N., Bertolino, E., Lin, Y. C., Laslo, P., Cheng, J. X., Murre, C., Singh, H., & Glass, C. K. (2010). Simple combinations of lineage-determining transcription factors prime cis-regulatory elements required for macrophage and B cell identities. Mol Cell, 38(4), 576–589. https://doi.org/10.1016/j.molcel.2010.05.004

33. Heuberger, J., & Birchmeier, W. (2010). Interplay of cadherin-mediated cell adhesion and canonical Wnt signaling. Cold Spring Harb Perspect Biol, 2(2), a002915. https://doi.org/10.1101/cshperspect.a002915

34. Hochane, M., van den Berg, P. R., Fan, X., Berenger-Currias, N., Adegeest, E., Bialecka, M., Nieveen, M., Menschaart, M., Chuva de Sousa Lopes, S. M., & Semrau, S. (2019). Single-cell transcriptomics reveals gene expression dynamics of human fetal kidney development. PLoS Biol, 17(2), e3000152. https://doi.org/10.1371/journal.pbio.3000152

35. Hodgkins, A., Farne, A., Perera, S., Grego, T., Parry-Smith, D. J., Skarnes, W. C., & Iyer, V. (2015). WGE: a CRISPR database for genome engineering. Bioinformatics, 31(18), 3078–3080. https://doi.org/10.1093/bioinformatics/btv308

36. Horster, M. F., Braun, G. S., & Huber, S. M. (1999). Embryonic renal epithelia: induction, nephrogenesis, and cell differentiation. Physiol Rev, 79(4), 1157–1191. https://doi.org/10.1152/physrev.1999.79.4.1157

37. Howe, L. R., Watanabe, O., Leonard, J., & Brown, A. M. (2003). Twist is up-regulated in response to Wnt1 and inhibits mouse mammary cell differentiation. Cancer Res, 63(8), 1906–1913. https://www.ncbi.nlm.nih.gov/pubmed/12702582

38. Huang, S. M., Mishina, Y. M., Liu, S., Cheung, A., Stegmeier, F., Michaud, G. A., Charlat, O., Wiellette, E., Zhang, Y., Wiessner, S., Hild, M., Shi, X., Wilson, C. J., Mickanin, C., Myer, V., Fazal, A., Tomlinson, R., Serluca, F., Shao, W., . . . Cong, F. (2009). Tankyrase inhibition stabilizes axin and antagonizes Wnt signalling. Nature, 461(7264), 614–620. https://doi.org/10.1038/nature08356

39. Ibar, C., Kirichenko, E., Keepers, B., Enners, E., Fleisch, K., & Irvine, K. D. (2018). Tension-dependent regulation of mammalian Hippo signaling through LIMD1. J Cell Sci, 131(5). https://doi.org/10.1242/jcs.214700

40. Kaku, Y., Taguchi, A., Tanigawa, S., Haque, F., Sakuma, T., Yamamoto, T., & Nishinakamura, R. (2017). PAX2 is dispensable for in vitro nephron formation from human induced pluripotent stem cells. Sci Rep, 7(1), 4554. https://doi.org/10.1038/s41598-017-04813-3

41. Kemler, R., Hierholzer, A., Kanzler, B., Kuppig, S., Hansen, K., Taketo, M. M., de Vries, W. N., Knowles, B. B., & Solter, D. (2004). Stabilization of beta-catenin in the mouse zygote leads to premature epithelial-mesenchymal transition in the epiblast. Development, 131(23), 5817–5824. https://doi.org/10.1242/dev.01458

42. Korsunsky, I., Millard, N., Fan, J., Slowikowski, K., Zhang, F., Wei, K., Baglaenko, Y., Brenner, M., Loh, P. R., & Raychaudhuri, S. (2019). Fast, sensitive and accurate integration of single-cell data with Harmony. Nat Methods, 16(12), 1289–1296. https://doi.org/10.1038/s41592-019-0619-0

43. Kumar, S. V., Er, P. X., Lawlor, K. T., Motazedian, A., Scurr, M., Ghobrial, I., Combes, A. N., Zappia, L., Oshlack, A., Stanley, E. G., & Little, M. H. (2019). Kidney micro-organoids in suspension culture as a scalable source of human pluripotent stem cell-derived kidney cells. Development, 146(5). https://doi.org/10.1242/dev.172361

44. Lindstrom, N. O., Guo, J., Kim, A. D., Tran, T., Guo, Q., De Sena Brandine, G., Ransick, A., Parvez, R. K., Thornton, M. E., Baskin, L., Grubbs, B., McMahon, J. A., Smith, A. D., & McMahon, A. P. (2018). Conserved and Divergent Features of Mesenchymal Progenitor Cell Types within the Cortical Nephrogenic Niche of the Human and Mouse Kidney. J Am Soc Nephrol, 29(3), 806–824. https://doi.org/10.1681/ASN.2017080890

45. Lindstrom, N. O., McMahon, J. A., Guo, J., Tran, T., Guo, Q., Rutledge, E., Parvez, R. K., Saribekyan, G., Schuler, R. E., Liao, C., Kim, A. D., Abdelhalim, A., Ruffins, S. W., Thornton, M. E., Baskin, L., Grubbs, B., Kesselman, C., & McMahon, A. P. (2018). Conserved and Divergent Features of Human and Mouse Kidney Organogenesis. J Am Soc Nephrol, 29(3), 785–805. https://doi.org/10.1681/ASN.2017080887

46. Lindstrom, N. O., Sealfon, R., Chen, X., Parvez, R. K., Ransick, A., De Sena Brandine, G., Guo, J., Hill, B., Tran, T., Kim, A. D., Zhou, J., Tadych, A., Watters, A., Wong, A., Lovero, E., Grubbs, B. H., Thornton, M. E., McMahon, J. A., Smith, A. D., . . . McMahon, A. P. (2021). Spatial transcriptional mapping of the human nephrogenic program. Dev Cell, 56(16), 2381–2398 e2386. https://doi.org/10.1016/j.devcel.2021.07.017

47. Lindstrom, N. O., Tran, T., Guo, J., Rutledge, E., Parvez, R. K., Thornton, M. E., Grubbs, B., McMahon, J. A., & McMahon, A. P. (2018). Conserved and Divergent Molecular and Anatomic Features of Human and Mouse Nephron Patterning. J Am Soc Nephrol, 29(3), 825–840. https://doi.org/10.1681/ASN.2017091036

48. Livak, K. J., & Schmittgen, T. D. (2001). Analysis of relative gene expression data using real-time quantitative PCR and the 2(-Delta Delta C(T)) Method. Methods, 25(4), 402–408. https://doi.org/10.1006/meth.2001.1262

49. Mah, S. P., Saueressig, H., Goulding, M., Kintner, C., & Dressler, G. R. (2000). Kidney development in cadherin-6 mutants: delayed mesenchyme-to-epithelial conversion and loss of nephrons. Dev Biol, 223(1), 38–53. https://doi.org/10.1006/dbio.2000.9738

50. Marable, S. S., Chung, E., & Park, J. S. (2020). Hnf4a Is Required for the Development of Cdh6-Expressing Progenitors into Proximal Tubules in the Mouse Kidney. J Am Soc Nephrol, 31(11), 2543–2558. https://doi.org/10.1681/ASN.2020020184

51. Marneros, A. G. (2020). AP-2beta/KCTD1 Control Distal Nephron Differentiation and Protect against Renal Fibrosis. Dev Cell, 54(3), 348–366 e345. https://doi.org/10.1016/j.devcel.2020.05.026

52. Massa, F., Garbay, S., Bouvier, R., Sugitani, Y., Noda, T., Gubler, M. C., Heidet, L., Pontoglio, M., & Fischer, E. (2013). Hepatocyte nuclear factor 1beta controls nephron tubular development. Development, 140(4), 886–896. https://doi.org/10.1242/dev.086546

53. McMahon, A. P. (2016). Development of the Mammalian Kidney. Curr Top Dev Biol, 117, 31–64. https://doi.org/10.1016/bs.ctdb.2015.10.010

54. McNeill, H., & Reginensi, A. (2017). Lats1/2 Regulate Yap/Taz to Control Nephron Progenitor Epithelialization and Inhibit Myofibroblast Formation. J Am Soc Nephrol, 28(3), 852–861. https://doi.org/10.1681/ASN.2016060611

55. Meeus, L., Gilbert, B., Rydlewski, C., Parma, J., Roussie, A. L., Abramowicz, M., Vilain, C., Christophe, D., Costagliola, S., & Vassart, G. (2004). Characterization of a novel loss of function mutation of PAX8 in a familial case of congenital hypothyroidism with in-place, normal-sized thyroid. J Clin Endocrinol Metab, 89(9), 4285–4291. https://doi.org/10.1210/jc.2004-0166

56. Naiman, N., Fujioka, K., Fujino, M., Valerius, M. T., Potter, S. S., McMahon, A. P., & Kobayashi, A. (2017). Repression of Interstitial Identity in Nephron Progenitor Cells by Pax2 Establishes the Nephron-Interstitium Boundary during Kidney Development. Dev Cell, 41(4), 349–365 e343. https://doi.org/10.1016/j.devcel.2017.04.022

57. Narlis, M., Grote, D., Gaitan, Y., Boualia, S. K., & Bouchard, M. (2007). Pax2 and pax8 regulate branching morphogenesis and nephron differentiation in the developing kidney. J Am Soc Nephrol, 18(4), 1121–1129. https://doi.org/10.1681/ASN.2006070739

58. Ng-Blichfeldt, J. P., & Röper, K. (2021). Mesenchymal-to-Epithelial Transitions in Development and Cancer. Methods Mol Biol, 2179, 43–62. https://doi.org/10.1007/978-1-0716-0779-4_7

59. Nishinakamura, R. (2019). Human kidney organoids: progress and remaining challenges. Nat Rev Nephrol, 15(10), 613–624. https://doi.org/10.1038/s41581-019-0176-x

60. Park, J. S., Ma, W., O’Brien, L. L., Chung, E., Guo, J. J., Cheng, J. G., Valerius, M. T., McMahon, J. A., Wong, W. H., & McMahon, A. P. (2012). Six2 and Wnt regulate self-renewal and commitment of nephron progenitors through shared gene regulatory networks. Dev Cell, 23(3), 637–651. https://doi.org/10.1016/j.devcel.2012.07.008

61. Park, J. S., Valerius, M. T., & McMahon, A. P. (2007). Wnt/beta-catenin signaling regulates nephron induction during mouse kidney development. Development, 134(13), 2533–2539. https://doi.org/10.1242/dev.006155

62. Patel, S. A., Hirosue, S., Rodrigues, P., Vojtasova, E., Richardson, E. K., Ge, J., Syafruddin, S. E., Speed, A., Papachristou, E. K., Baker, D., Clarke, D., Purvis, S., Wesolowski, L., Dyas, A., Castillon, L., Caraffini, V., Bihary, D., Yong, C., Harrison, D. J., . . . Vanharanta, S. (2022). The renal lineage factor PAX8 controls oncogenic signalling in kidney cancer. Nature, 606(7916), 999–1006. https://doi.org/10.1038/s41586-022-04809-8

63. Pei, D., Shu, X., Gassama-Diagne, A., & Thiery, J. P. (2019). Mesenchymal-epithelial transition in development and reprogramming. Nat Cell Biol, 21(1), 44–53. https://doi.org/10.1038/s41556-018-0195-z

64. Phelps, D. E., & Dressler, G. R. (1996). Identification of novel Pax-2 binding sites by chromatin precipitation. J Biol Chem, 271(14), 7978–7985. https://doi.org/10.1074/jbc.271.14.7978

65. Plachov, D., Chowdhury, K., Walther, C., Simon, D., Guenet, J. L., & Gruss, P. (1990). Pax8, a murine paired box gene expressed in the developing excretory system and thyroid gland. Development, 110(2), 643–651. https://doi.org/10.1242/dev.110.2.643

66. Raudvere, U., Kolberg, L., Kuzmin, I., Arak, T., Adler, P., Peterson, H., & Vilo, J. (2019). g:Profiler: a web server for functional enrichment analysis and conversions of gene lists (2019 update). Nucleic Acids Res, 47(W1), W191–W198. https://doi.org/10.1093/nar/gkz369

67. Rauskolb, C., Sun, S., Sun, G., Pan, Y., & Irvine, K. D. (2014). Cytoskeletal tension inhibits Hippo signaling through an Ajuba-Warts complex. Cell, 158(1), 143–156. https://doi.org/10.1016/j.cell.2014.05.035

68. Rodriguez-Boulan, E., & Macara, I. G. (2014). Organization and execution of the epithelial polarity programme. Nat Rev Mol Cell Biol, 15(4), 225–242. https://doi.org/10.1038/nrm3775

69. Rothenpieler, U. W., & Dressler, G. R. (1993). Pax-2 is required for mesenchyme-to-epithelium conversion during kidney development. Development, 119(3), 711–720. https://doi.org/10.1242/dev.119.3.711

70. Schep, A. N., Wu, B., Buenrostro, J. D., & Greenleaf, W. J. (2017). chromVAR: inferring transcription-factor-associated accessibility from single-cell epigenomic data. Nat Methods, 14(10), 975–978. https://doi.org/10.1038/nmeth.4401

71. Schmidt-Ott, K. M., Masckauchan, T. N., Chen, X., Hirsh, B. J., Sarkar, A., Yang, J., Paragas, N., Wallace, V. A., Dufort, D., Pavlidis, P., Jagla, B., Kitajewski, J., & Barasch, J. (2007). beta-catenin/TCF/Lef controls a differentiation-associated transcriptional program in renal epithelial progenitors. Development, 134(17), 3177–3190. https://doi.org/10.1242/dev.006544

72. Schreibing, F., & Kramann, R. (2022). Mapping the human kidney using single-cell genomics. Nat Rev Nephrol, 18(6), 347–360. https://doi.org/10.1038/s41581-022-00553-4

73. Schunk, S. J., Floege, J., Fliser, D., & Speer, T. (2021). WNT-beta-catenin signalling - a versatile player in kidney injury and repair. Nat Rev Nephrol, 17(3), 172–184. https://doi.org/10.1038/s41581-020-00343-w

74. Severin, J., Lizio, M., Harshbarger, J., Kawaji, H., Daub, C. O., Hayashizaki, Y., Consortium, F., Bertin, N., & Forrest, A. R. (2014). Interactive visualization and analysis of large-scale sequencing datasets using ZENBU. Nat Biotechnol, 32(3), 217–219. https://doi.org/10.1038/nbt.2840

75. Stewart, B. J., Ferdinand, J. R., Young, M. D., Mitchell, T. J., Loudon, K. W., Riding, A. M., Richoz, N., Frazer, G. L., Staniforth, J. U. L., Vieira Braga, F. A., Botting, R. A., Popescu, D. M., Vento-Tormo, R., Stephenson, E., Cagan, A., Farndon, S. J., Polanski, K., Efremova, M., Green, K., . . . Clatworthy, M.R. (2019). Spatiotemporal immune zonation of the human kidney. Science, 365(6460), 1461–1466. https://doi.org/10.1126/science.aat5031

76. Stewart, B. J., Fergie, M., Young, M., Jones, C., Sachdeva, A., Blain, A. E., Bacon, C. M., Rand, V., Ferdinand, J. R., James, K. R., Mahbubani, K. T., Hook, C. E., Jonas, N., Coleman, N., Saeb-Parsy, K., Collin, M., Clatworthy, M., Behjati, S., & Carey, C. D. (2023). Spatial and molecular profiling of the mononuclear phagocyte network in Classic Hodgkin lymphoma. Blood. https://doi.org/10.1182/blood.2022015575

77. Stuart, T., Butler, A., Hoffman, P., Hafemeister, C., Papalexi, E., Mauck, W. M., 3rd, Hao, Y., Stoeckius, M., Smibert, P., & Satija, R. (2019). Comprehensive Integration of Single-Cell Data. Cell, 177(7), 1888–1902 e1821. https://doi.org/10.1016/j.cell.2019.05.031

78. Takasato, M., Er, P. X., Chiu, H. S., Maier, B., Baillie, G. J., Ferguson, C., Parton, R. G., Wolvetang, E. J., Roost, M. S., Chuva de Sousa Lopes, S. M., & Little, M.H. (2015). Kidney organoids from human iPS cells contain multiple lineages and model human nephrogenesis. Nature, 526(7574), 564–568. https://doi.org/10.1038/nature15695

79. Tan, G., & Lenhard, B. (2016). TFBSTools: an R/bioconductor package for transcription factor binding site analysis. Bioinformatics, 32(10), 1555–1556. https://doi.org/10.1093/bioinformatics/btw024

80. Tang, W. W. C., Castillo-Venzor, A., Gruhn, W. H., Kobayashi, T., Penfold, C. A., Morgan, M. D., Sun, D., Irie, N., & Surani, M. A. (2022). Sequential enhancer state remodelling defines human germline competence and specification. Nat Cell Biol, 24(4), 448–460. https://doi.org/10.1038/s41556-022-00878-z

81. Torres, M., Gomez-Pardo, E., Dressler, G. R., & Gruss, P. (1995). Pax-2 controls multiple steps of urogenital development. Development, 121(12), 4057–4065. https://doi.org/10.1242/dev.121.12.4057

82. Vallin, J., Thuret, R., Giacomello, E., Faraldo, M. M., Thiery, J. P., & Broders, F. (2001). Cloning and characterization of three Xenopus slug promoters reveal direct regulation by Lef/beta-catenin signaling. J Biol Chem, 276(32), 30350–30358. https://doi.org/10.1074/jbc.M103167200

83. van Dijk, D., Sharma, R., Nainys, J., Yim, K., Kathail, P., Carr, A. J., Burdziak, C., Moon, K. R., Chaffer, C. L., Pattabiraman, D., Bierie, B., Mazutis, L., Wolf, G., Krishnaswamy, S., & Pe’er, D. (2018). Recovering Gene Interactions from Single-Cell Data Using Data Diffusion. Cell, 174(3), 716–729 e727. https://doi.org/10.1016/j.cell.2018.05.061

84. Vierstra, J., Lazar, J., Sandstrom, R., Halow, J., Lee, K., Bates, D., Diegel, M., Dunn, D., Neri, F., Haugen, E., Rynes, E., Reynolds, A., Nelson, J., Johnson, A., Frerker, M., Buckley, M., Kaul, R., Meuleman, W., & Stamatoyannopoulos, J. A. (2020). Global reference mapping of human transcription factor footprints. Nature, 583(7818), 729–736. https://doi.org/10.1038/s41586-020-2528-x

85. Wickham, H. (2016). ggplot2: Elegant Graphics for Data Analysis (Springer-Verlag).

86. Yamashita, K., Sato, A., Asashima, M., Wang, P. C., & Nishinakamura, R. (2007). Mouse homolog of SALL1, a causative gene for Townes-Brocks syndrome, binds to A/T-rich sequences in pericentric heterochromatin via its C-terminal zinc finger domains. Genes Cells, 12(2), 171–182. https://doi.org/10.1111/j.1365-2443.2007.01042.x

87. Yook, J. I., Li, X. Y., Ota, I., Fearon, E. R., & Weiss, S. J. (2005). Wnt-dependent regulation of the E-cadherin repressor snail. J Biol Chem, 280(12), 11740–11748. https://doi.org/10.1074/jbc.M413878200

88. Yook, J. I., Li, X. Y., Ota, I., Hu, C., Kim, H. S., Kim, N. H., Cha, S. Y., Ryu, J. K., Choi, Y. J., Kim, J., Fearon, E. R., & Weiss, S. J. (2006). A Wnt-Axin2-GSK3beta cascade regulates Snail1 activity in breast cancer cells. Nat Cell Biol, 8(12), 1398–1406. https://doi.org/10.1038/ncb1508

89. Zhang, Y., Liu, T., Meyer, C. A., Eeckhoute, J., Johnson, D. S., Bernstein, B. E., Nusbaum, C., Myers, R. M., Brown, M., Li, W., & Liu, X. S. (2008). Model-based analysis of ChIP-Seq (MACS). Genome Biol, 9(9), R137. https://doi.org/10.1186/gb-2008-9-9-r137

90. Zhao, B., Ye, X., Yu, J., Li, L., Li, W., Li, S., Yu, J., Lin, J. D., Wang, C. Y., Chinnaiyan, A. M., Lai, Z. C., & Guan, K. L. (2008). TEAD mediates YAP-dependent gene induction and growth control. Genes Dev, 22(14), 1962–1971. https://doi.org/10.1101/gad.1664408

